# Synthetic Immunological Niche Reveals Early Immune Dysregulation and Stratifies Therapeutic Response in Type 1 Diabetes

**DOI:** 10.64898/2026.03.06.710219

**Authors:** Jyotirmoy Roy, Yifei Jiang, Runbo Mao, Jessica L. King, Amod Talekar, Lillian Holman, Haoxuan Zeng, Xin Luo, Peter Sajjakulnukit, Brianna Ha, Kai Liu, Elizabeth J. Bealer, Laila M. Rad, Shahzad Sohail, Antonio Holmes, Bryan Wonski, Kathryn Kang, Dominik Awad, Aaron H. Morris, Costas A. Lyssiotis, Jie Liu, Lonnie D. Shea

## Abstract

In Type 1 Diabetes (T1D), disease onset and response to immunotherapy vary widely among individuals, reflecting heterogeneous stage-specific immune dysregulation that remains undefined. To investigate this heterogeneity, we engineered a microporous polycaprolactone scaffold that forms a synthetic immunological niche (IN) upon subcutaneous implantation, enabling in vivo capture of systemic immune dysregulation. In non-obese diabetic (NOD) mice, longitudinal transcriptomic profiling of IN-infiltrating cells identified early-stage genes relatively enriched for myeloid cells, followed by progressive increases in T cell-associated dysregulation at later stages that distinguished T1D progressors from non-progressors. We derived an early-stage IN-based T1D gene signature capturing immune alterations. The signature stratified NOD progressors from non-progressors as early as 6 weeks of age and was conserved across human T1D datasets, distinguishing T1D from non-diabetic individuals in spleen and pancreatic lymph node samples, but not peripheral blood. Signature-based stratification further revealed enrichment of macrophage-associated TNF-α pathways in NOD progressors, validated in human T1D islets. Given heterogeneous response to anti-TNF-α therapy, IN profiling identified resistance-associated mechanisms and enabled derivation of a pathway score that prospectively distinguished treatment-sensitive from resistant mice prior to therapy, establishing the IN as a minimally invasive platform for detecting stage-wise immune dysregulation and stratifying immunotherapy response in T1D.

## Introduction

Type 1 diabetes (T1D) is an autoimmune disease characterized by immune cell infiltration of the pancreatic islets, leading to the destruction of insulin-producing β cells (1). T1D progression is marked by substantial immunological heterogeneity across individuals, with variability in immune cell composition, activation states, and inflammatory programs shaping distinct disease trajectories (2). While T1D has been associated with genetic susceptibility, particularly mutations in the class II human leukocyte antigen (HLA) region, only 5% or fewer with these risk alleles in the general population progress to overt hyperglycemia (3–6). Blood-based autoantibodies, another clinical marker for T1D, also represent relatively late manifestations of the autoimmune T1D process, occurring downstream of early innate immune dysregulation (7–9). Importantly, not all autoantibody-positive individuals ultimately progress to overt diabetes, limiting their predictive precision in early stages (10). As a result, immunotherapies are administered in T1D patients at the late stages, when full-fledged adaptive immune dysregulation has begun and a substantial β cell loss has occurred, resulting in sub-optimal outcomes.

The limited clinical efficacy outcomes of immunotherapies are reflected in the heterogeneous therapeutic responses observed across individuals (11). Teplizumab, an anti-CD3 monoclonal antibody administered during the later stage of T1D, can only delay the clinical onset by a median of about two years (11). Similarly, Baricitinib is a JAK inhibitor that can preserve β-cell function in newly diagnosed stage 3 diabetes, characterized by onset of hyperglycemia along with presence of multiple autoantibodies. However, this therapy only slows T1D progression and as with most immunotherapies, is not effective in all patients (12). The variable efficacy of these immunotherapies in T1D reflects the heterogeneity and stage-specific nature of immune dysregulation, which complicates patient stratification and therapeutic timing. These limitations in understanding immune heterogeneity in T1D highlight the need for an in-vivo diagnostic platform that can provide insight into stage-wise immune dysregulation preceding substantial β cell destruction.

In this report, we apply a microporous polycaprolactone (PCL) scaffold that, when implanted subcutaneously establishes a synthetic immunological niche (IN) in vivo, and captures stage-wise immune dysregulation preceding overt diabetes onset. The porous architecture of the IN promotes host immune cell infiltration and vascularization, allowing recruitment of immune populations. Previous studies from our lab have demonstrated that this IN microenvironment reflects tissue-specific immune dynamics and can identify signatures related to disease progression in multiple disease contexts, including triple-negative breast cancer, organ transplantation, and autoimmune diseases such as multiple sclerosis (13–18). In T1D, our previous work has shown that the IN transcriptomic based scoring can monitor disease progression in the Non-obese diabetic (NOD) model (19). Herein, we apply the IN to reveal the underlying immune inflammatory programs that distinguish T1D progressors from non-progressors in NOD mice over the stages of disease and validate that these programs are conserved in human T1D. We identify a macrophage-associated enriched TNF-α inflammatory profile in T1D progressors, which is also validated in human T1D islets. Therapeutic targeting with anti-TNF-α revealed heterogeneity in treatment response, which was captured by the IN and quantified through a pathway response score that distinguished treatment-sensitive from resistant individuals. Collectively, these findings establish the synthetic IN as a minimally invasive platform capable of detecting stage-wise immune inflammatory perturbations, identifying actionable immunotherapy targets and guiding personalized immunotherapy strategies in T1D.

## Results

### Immunological niche captures early immune and inflammatory dysregulation in T1D

We initially aimed to identify stage-associated immune alterations underlying T1D pathogenesis by performing longitudinal analysis of the immune composition and related transcriptomic profile of the IN. Microporous polycaprolactone (PCL) scaffolds were implanted subcutaneously in female NOD mice at 4 weeks of age (**Figure 1a**). Due to their microporous structure, these scaffolds became vascularized and recruited immune cells, forming a synthetic IN. Building on our previous work which showed that the IN microenvironment contains immune cell types representative of the NOD pancreas, we sought to define how these cell types and related genes change across disease stage by explanting the IN with replacement at early (6 weeks), intermediate (12-14 weeks), and late (17-20 weeks) stages for immunophenotyping and transcriptomic analysis (20).

**Figure 1.**
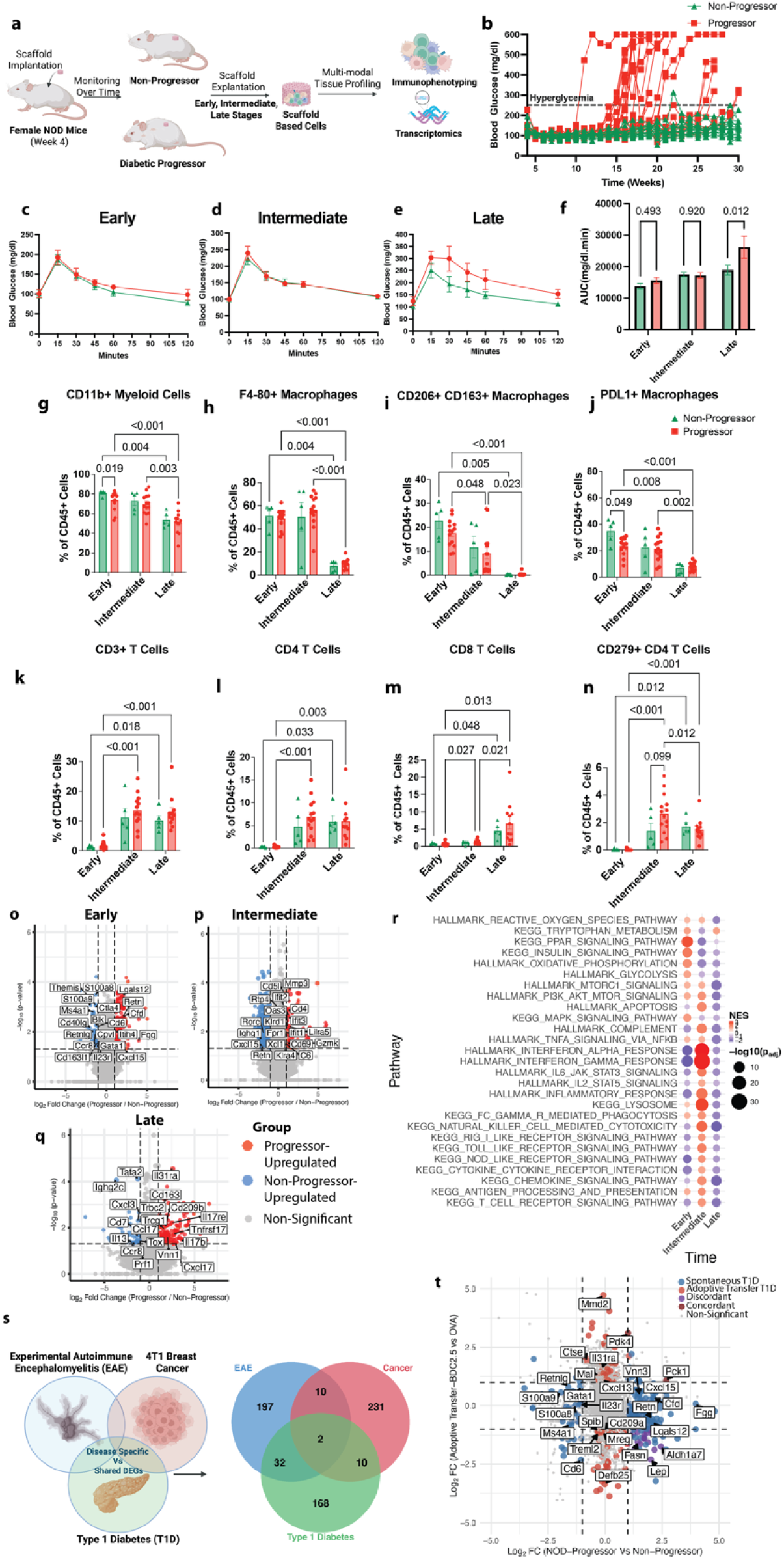
Immunological niche captures early immune and inflammatory dysregulation specific to T1D. **a,** Experimental design: Female Non-Obese Diabetic (NOD) mice were implanted with the scaffolds at 4 weeks of age. Scaffold based immunological niches (INs) were explanted at early (∼6 weeks), intermediate (∼11-15 weeks), and late (∼16-20 weeks) stages to characterize immune and transcriptomic changes in diabetic progressors (blood glucose ≥250 mg/dl) versus non-progressors (normoglycemic). **b,** Blood glucose levels per mice over time: By 30 weeks, the progressors mice developed hyperglycemia (blood glucose ≥250mg/dl), while the non-progressors remained normoglycemic). **c-f,** Intraperitoneal Glucose Tolerance Test (IPGTT) in progressor (n-5) vs non-progressor (n=4) at **c,** early, **d,** intermediate, **e,** late stage and **f,** AUC for progressor vs non-progressor across each stage. Statistical analysis was performed using multiple unpaired t-tests with single pooled variance. Data are shown as mean ± SEM. **g–n,** Immune cell composition in the IN analyzed by flow cytometry as fraction of CD45+ immune cells: **g,** Myeloid cells **h,** macrophages **i,** M2-like macrophages (CD206⁺CD163⁺) **j,** PD-L1⁺ **k,** T cells, **l,** CD4⁺ T cells, **m,** CD8⁺ T cells, **n,** PD-1⁺ CD4⁺ T cells as a fraction of total immune cells. Statistical analysis was performed using a mixed-effects model with the Geisser-Greenhouse correction, followed by Tukey’s multiple-comparison test, with individual variances computed for each comparison. Data are shown as mean ± SEM (n = 5-14 per group). **o–q,** Differential Gene Expression Analysis of IN transcriptomics across disease stages: Volcano plots of DEGs (p ≤ 0.05, |log₂FC| ≥ 1) in progressors vs non-progressors at **o,** early (n = 16-24 per group), **p,** intermediate (n = 10 per group), and **q,** late (n = 4-10 per group) stages. Axes were constrained for visualization (log₂FC: -8 to 8; -log_10_(p): 0 to 6). **r,** Genset enrichment analysis across disease stages: Dotplot of selected enriched pathways in progressor vs non-progressor (FDR ≤ 0.1, |NES| ≥ 1 in at least one stage). **s,** Venn Diagrams showing limited overlap between of IN based early stage T1D DEGs with IN based DEGs from 4T1 Breast cancer and Experimental Autoimmune Encephalomyelitis model **t,** Double Volcano Plot showing fold changes of IN based early stage DEGs in spontaneous NOD T1D model compared to IN DEGs from Adoptive Transfer T1D model. Genes colored show differential expression (p ≤ 0.05, |log₂FC| ≥ 1). Axes were constrained to ±5 log₂FC for visualization.

Progressor mice were defined as those that developed hyperglycemia (blood glucose ≥250 mg/dL), whereas non-progressors maintained normoglycemia through 30 weeks of age (**Figure 1b, Supplementary Table 1**). Notably, conventional systemic readouts like intraperitoneal glucose tolerance testing (IPGTT) failed to distinguish T1D progressors from non-progressors until the late stage (**Figure 1c-f, Supplementary Table 1**).

The limited early predictive power of these conventional systemic readouts motivated an examination of differences within the IN. The immune cell composition of the niche microenvironment was characterized by spectral immunophenotyping (**Figure 1g-n, Supplementary Figure 1, Supplementary Table 2**). At the early and intermediate stages, the IN immune compartment was dominated by myeloid cells, particularly macrophages, in both progressors and non-progressors (**Figure 1g–h**). The proportion of anti-inflammatory M2-like macrophages (CD163^+^CD206⁺) progressively declined in both groups, indicating a general shift toward a pro-inflammatory state (**Figure 1i**). The proportion of PD-L1⁺ macrophages was slightly higher in non-progressors at early stages than progressors, indicating inhibitory function related differences (**Figure 1j**). We next characterized the lymphoid compartment and identified that the proportion of T cells increased within the IN immune microenvironment from the intermediate stage onwards in both groups (**Figure 1k**). The CD4⁺ T cell proportion significantly increased by the intermediate stage, while CD8⁺ T cells had a pronounced increase at the late stage of T1D (**Figure 1l-m**). Notably, exhausted CD279+ CD4⁺ T cells were modestly increased in the T1D progressors vs non-progressors at the intermediate stage, suggestive of increased activation (**Figure 1n**). Since differences in immune cell proportions between the T1D progressors and non-progressors were modest, we next interrogated transcriptional phenotypes of these cells to uncover functional phenotypic distinctions not captured at the level of cellular abundance.

Differential gene expression analysis of bulk RNA sequencing data from IN-derived cells was performed to identify stage-wise transcriptional differences between progressor and non-progressor NOD mice (**Figure 1o–q**). We identified 212, 600, and 242 differentially expressed genes (DEGs; *p* < 0.05 and |log₂FC| > 1) at the early, intermediate, and late stages, respectively (**Supplementary Table 3)**. At the early stage, progressors exhibited a transcriptional program dominated by innate myeloid dysregulation, marked by upregulation of genes like *Cxcl15, Cfd, Itih4, Vnn3, Fgg, Lglas12,* and *Retn* and downregulation of *S100a8, S100a9, Cd163l1 and Cpvl.* Alterations in some lymphoid associated genes such as *Il23r, Cpvl, Ccr8, Cd40lg, Ms4a, Cd6,* and *Themis* were also present. Together these changes indicate a dysregulated myeloid compartment with some differences in adaptive immune signaling in progressors vs non-progressors at the early stage (**Figure 1o**). At the intermediate stage, there was increased representation of adaptive immune dysregulation. Progressors exhibited upregulation of interferon-stimulated genes like *Ifit1*, *Ifit2*, *Ifit3*, *Oas3* and *Rtp4*, cytotoxic and activation-associated genes like *Gzmk*, *Klra5*, *Klrd1*, *Cd69*, *Cd7* and Cd*4,* and downregulation of *Rorc and Ighg1*. Some innate immune related genes like *C6*, *Lilra5*, *Cxcl15, Retn* and *Cd5l* were also differentially expressed (**Figure 1p**). At the late stage, progressors exhibited upregulation of inflammatory and immune activation markers such as *Tnfrsf17, Cd209b, Il17re, Il31ra,* and *Cxcl1*7 (**Figure 1q**).

To extend the transcriptomic analysis beyond individual genes, Gene Set Enrichment Analysis (GSEA) was performed to identify the related differentially regulated pathways between T1D progressor and non-progressor across the disease stages (**Figure 1r**, **Supplementary Table 3**). Consistent with gene level findings, progressors demonstrated higher enrichment of inflammatory immune pathways associated with TNF-α via NF-κ,β signaling, IL-6 JAK-STAT signaling, IFN-γ/α response, reactive oxygen species (ROS) production, T cell receptor signaling and antigen from early to intermediate stages, reflecting increasing immune inflammatory differences between the progressors vs non-progressors. Interestingly, at the late stage, many of these pathways displayed relative downregulation in progressors, possibly hinting at subsiding of inflammation after substantial β cell destruction. Given our focus on preclinical immune divergence and early immune dysregulation, we focused on the early stage for subsequent analysis.

We next evaluated the specificity of the IN-based early stage DEGs to T1D pathogenesis. We compared the early stage T1D DEGs with those from other inflammatory disease models like 4T1 breast cancer and progressive Experimental Autoimmune Encephalomyelitis (EAE) (21) (**Figure 1s**). Notably, the IN-based DEGs in diseased versus healthy condition across these three disease models were largely disease-specific, with limited overlap. Of the 212 T1D DEGs, only 34 overlapped with those observed in the autoimmune disease EAE, with just 12 genes shared with 4T1 breast cancer, while 168 DEGs were specific to T1D. (**Figure 1s**). Similarly, the pathway enrichment analysis using GSEA revealed that the IN captured distinct enrichment patterns in signaling pathways like PPAR, interferon, and mTORC1 signaling across the diseases (**Supplementary Figure 2a, Supplementary Table 4)**.

Having established disease specificity, we compared early stage DEGs from the spontaneous NOD model with those from the T cell adoptive-transfer model in NOD.scid mice and found limited overlap, reflecting distinct disease related programs across these two settings (20). DEGs in the adoptive transfer model consisted of adaptive immune inflammation genes like *Il31ra* and *Spib* and were largely distinct from those observed in the early stage of the spontaneous NOD model (**Figure 1t**). The associated pathways also highlighted a predominant upregulation of adaptive immune cell associated gene programs like IFN-γ (**Supplementary Figure 2b, Supplementary Table 4**). Together, these findings indicate that the IN not only captures T1D-related transcriptional changes but also distinguishes between distinct immunopathogenic contexts, differentiating early stages of spontaneous T1D from acute, T cell-mediated diabetes.

The transcriptional divergence between progressors and non-progressors led to an investigation of the metabolic landscape of the IN microenvironment to determine whether similar differences were evident (**Supplementary Table 5)**. Lipidomic profiling at the early stage showed minimal variation in lipid composition between groups (**Supplementary Figure 3a**). Broader metabolomic profiling demonstrated a more time-dependent shift with metabolites related to amino acid metabolism, the tricarboxylic acid (TCA) cycle, and glycolysis upregulated at early stage indicating highly metabolically active states of the cells. Nucleotide-related metabolites increased at the intermediate and late stage, indicating higher cell proliferation (**Supplementary Figure 3b**). However, no significant differences between the progressors vs non-progressor were observed at the early stages. Given that these metabolic alterations were secondary to the more pronounced immune transcriptional divergence between progressors vs non-progressors, subsequent analyses focused on differences at the transcriptomic level.

### IN-based immune DEGs are reflective of immune microenvironment of NOD pancreas and vital human T1D tissues

We next tested the hypothesis that the IN based immune related DEGs reflect the immune microenvironment of T1D relevant tissues. A representative immune subset of early and intermediate stage IN DEGs were mapped onto single cell-RNA sequencing datasets from pancreas resident immune cells in NOD mice as well as pancreatic lymph node (pLN), spleen, and PBMCs from T1D patients and non-diabetic controls (**Figure 2a)**.

**Figure 2.**
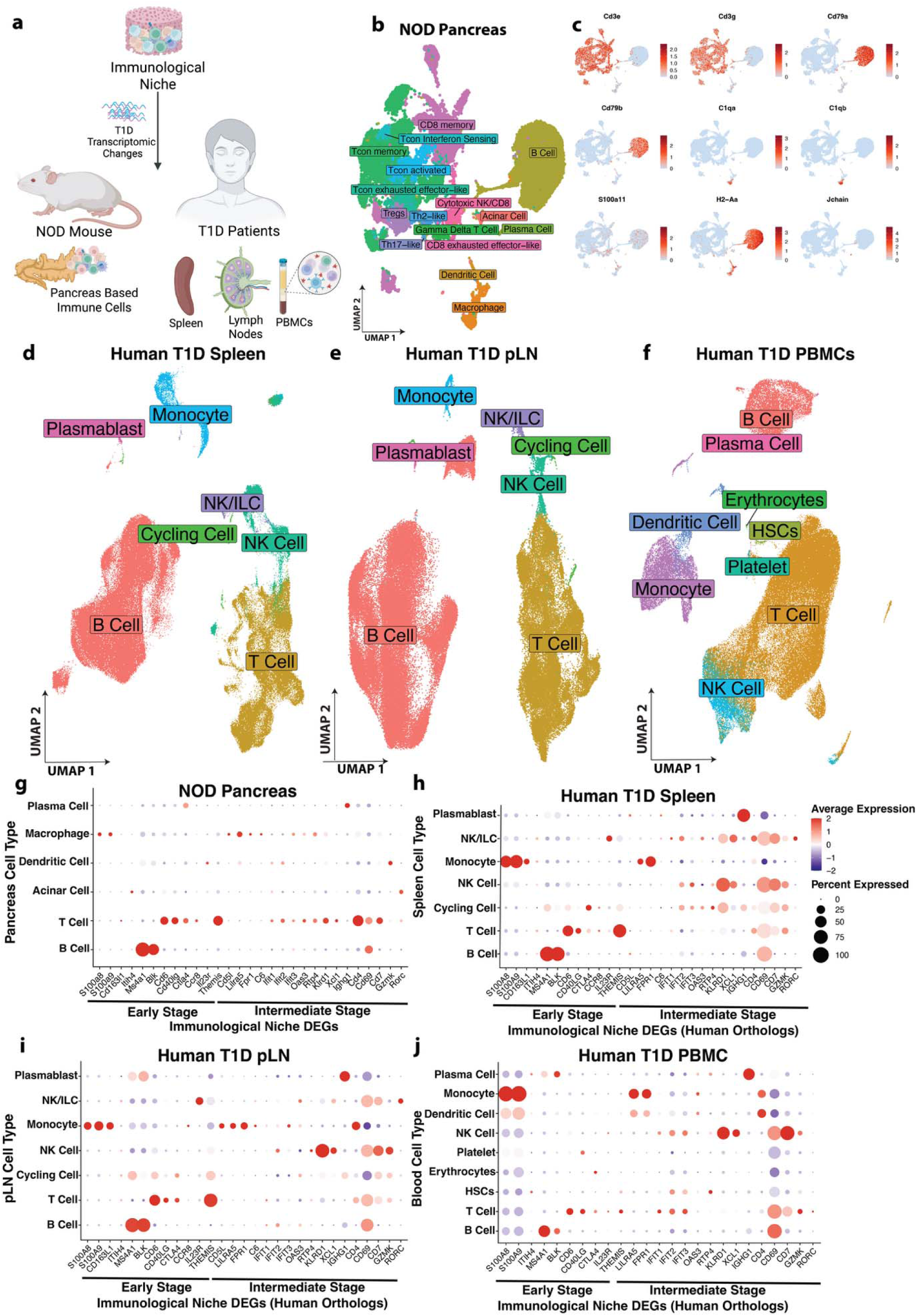
IN-based immune DEGs are reflective of immune microenvironment of NOD Pancreas and Vital Human T1D tissues. **a,** IN based early and intermediate stage DEGs were mapped to immune cells from NOD pancreas and spleen, pancreatic lymph node (pLN) and PBMCs from human T1D individuals and healthy control. **b**, UMAP of immune cells sorted from NOD pancreas at early and intermediate stages. **c,** Feature plot of canonical genes used for identifying cell types in NOD pancreas. **d-f,** UMAP showing cell types from human T1D individuals and healthy control in **d**, spleen **e**, pLN and **f**, PBMCs, **g-j**. Dotplot showing scaled average expression and percentage expression of IN based early and intermediate stage DEGs and their human orthologs in **g**, NOD pancreas, and **h**, human spleen, **i**, pLN, **j**, PBMCs from T1D individuals and non-diabetic control.

In the NOD model, pancreatic immune cells from mice were isolated by fluorescence-activated cell sorting (FACS) at early (6 weeks) and intermediate (12 weeks) stages and profiled by single-cell RNA sequencing (**Supplementary Figure 4)**. Major immune cell types were identified using a graph-based clustering approach and Uniform Manifold Approximation and Projection (UMAP) dimensionality reduction implemented in the Seurat package (22–25) (**Figure 2b**). Cell identities were assigned using canonical marker genes (**Figure 2c, Supplementary Figure 5).** For IN-DEG mapping purposes, T cell subsets were grouped broadly as T cells. Human spleen, pancreatic lymph node (pLN), and PBMC single-cell datasets were processed and annotated using dataset-specific clustering and cell-type annotation strategies as described in the Methods (Human Validation), enabling identification of key innate and adaptive immune populations, including monocytes, dendritic cells, NK cells, T cells, and B cells (**Figure 2d-e**).

Mapping of IN DEGs demonstrated representation across major immune compartments across vital tissues in NOD and T1D patients. In the NOD pancreas, early stage DEGs like *S100a8* and *S100a9* were enriched in the macrophage populations whereas genes like *Itih4*, *Ms4a1*, *Blk, Cd6, Cd40lg, Ctla4, Ccr8* and *Themis* were enriched within the B and T cell compartment (**Figure 2g**). A similar distribution pattern was observed in human spleen, pLN and PBMCs, where the human orthologs mapped to corresponding innate and adaptive immune populations (**Figure 2h-j**). At the intermediate stage, innate immune associated genes like *Cd5l*, *Lilra5,* Fpr*1,* and *C6* were mapped predominantly to pancreas macrophages in NOD, while their human orthologs were detected in monocytes from human tissues (**Figure 2g-j**). In parallel interferon-and cytotoxic-associated genes like ifit*1*, *Ifit2*, *Ifit3, Oas3, Rtp4, Klrd1, Xcl1, Cd4, Cd69, Cd7, Gzmk,* and *Rorc* were enriched in the T cell population in both NOD mouse pancreas and the human tissues (**Figure 2g-j**). These findings indicate that the IN-derived early and intermediate immune DEGs are reflective of myeloid and lymphoid populations in disease relevant tissues.

Notably, some of the IN immune DEGs, including *Cd163l1*, *Ccr8*, *Cd5l*, and *C6*, were found in the NOD pancreas, human T1D spleen and pLN but not prominently detected in peripheral blood, highlighting the IN’s ability to capture tissue-associated immune programs that may not be detectable through circulating biomarkers.

### Tissue-associated IN-based T1D gene signature stratifies early disease progression and is conserved across human tissues

We next aimed to identify an early-stage IN-derived T1D gene signature that captures disease-relevant, tissue-associated changes conserved in human T1D pathology, and serves as a predictive marker of disease progression. A discriminative gene panel was derived from early-stage transcriptomics of the IN by applying supervised partial least squares discriminant analysis (PLS-DA) to select a T1D gene signature comprising the top 100 discriminative genes (**Figure 3a, Supplementary Table 6)**. This 100 gene panel distinguished progressors from non-progressors at the early stage, indicating that coordinated gene programs, rather than individual DEGs, better capture early divergence in disease trajectory (**Figure 3a, b**). Biologically, the T1D gene signature encompassed multiple immune and cellular programs. These included innate immunity related genes like *Cfi*, *Cpvl, Psme3, Tm9sf4,* and *Tnfaip8l1* along with T cell-linked immune modulators and checkpoint-associated genes like *Cd40lg*, *Ccr8*, *Pdcd1*. In addition, transcriptional and chromatin regulators like *Sp8*, *Yy2*, *Ncor1*, and *Zfp70*3, and trafficking or protein turnover-related factors like *Rab3c*, *Snx3* and *Nedd4l* were also present. These various biological programs highlight that an IN-based T1D gene signature is indicative of immune inflammatory signaling and related transcriptional remodeling preceding overt T1D. This T1D gene signature was reflective of the innate and adaptive immune compartments in the NOD pancreas, like the immune DEGs (**Supplementary Figure 6)**.

**Figure 3:**
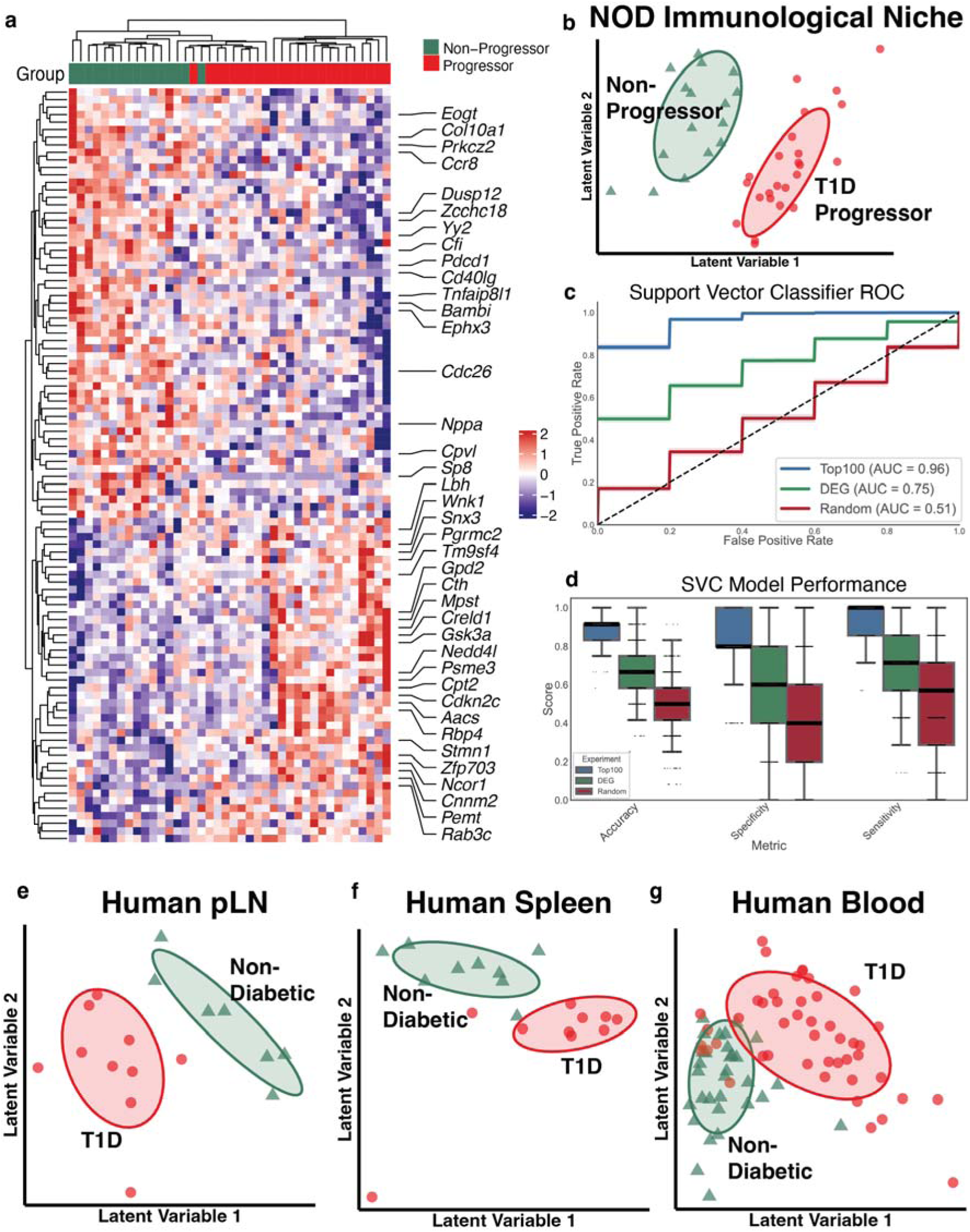
Tissue-Associated IN-Based T1D Gene Signature Stratifies Early Disease Progression and Is Conserved Across Human Tissues. **a,** Heatmap of IN-based T1D 100 gene signature identified using Partial Least Square Discriminatory Analysis (PLS-DA) at early stage in progressors vs non-progressors. **b,** PLS-DA score plot showing distinct clustering of NOD progressors and non-progressors using the IN-based T1D gene signature at early stage. **c,** AUC**-**ROC of SVC model trained using the IN-based T1D gene signature vs early stage DEGs and random 100 genes **e,** Box plots showing the performance of the SVC model in terms of accuracy, specificity and sensitivity for the IN-based T1D gene signature vs early stage DEGs and random 100 genes. **e-g,** PLS-DA score plots were used to assess separation between T1D and non-diabetic (ND) individuals based on human orthologs of the IN-derived T1D gene signature. These genes were projected onto pseudo-bulked single-cell datasets from human tissues. Clear separation between T1D and ND samples was observed in the **e**, pancreatic lymph node (pLN; T1D = 10, ND = 7) and **f**, spleen (T1D = 10, ND = 9), **g**, whereas peripheral blood (T1D = 46, ND = 31) exhibited partial overlap between groups. Ellipses indicate the one standard deviation (∼68%) multivariate normal contour for each group.

The predictive capacity of the T1D gene signature was quantitatively evaluated by training a Support Vector Classifier (SVC), implementing hyperparameter optimization and 1,000 bootstrapped iterations to ensure model robustness. The classifier demonstrated strong predictive performance and accurately predicted progressor vs non-progressor status at early stage, achieving an AUC-ROC of 0.96 and mean accuracy of 0.89, specificity of 0.81 and sensitivity of 0.95 (**Figure 3c, d)**. In contrast, identically parameterized SVC models trained on either full set of early stage 212 DEGs or randomly selected 100-gene panels yielded substantially reduced performance (Early Stage DEGs: AUC-ROC 0.75, accuracy 0.67, sensitivity 0.69, specificity 0.64; Random 100 genes: AUC-ROC 0.51, accuracy 0.49, sensitivity 0.49, specificity 0.49).

The human orthologs of the IN derived T1D gene signature were mapped onto pseudo-bulked data from spleen, pancreatic lymph node and PBMCs of individuals with T1D vs non-diabetic controls. Using PLS-DA for supervised dimensionality reduction, we assessed whether this signature could discriminate disease status in these human tissues. In both spleen and pLN, 93 of the 100 signature genes were detected, and samples segregated by disease status in PLS-DA space, demonstrating conservation of the IN-derived transcriptional programs in human T1D tissues (**Figure 3e, f**). In contrast, only 73 signature genes were present in PBMC dataset, and partial overlap between T1D and control samples was observed in PLS-DA projections indicating reduced discriminatory resolution in circulation compared to tissues (**Figure 3g**).

Collectively, these findings demonstrate that the IN-derived T1D gene signature captures disease-relevant immune programs that are preserved across species and more prominently reflected within tissue-based cellular phenotypes, highlighting its potential to detect early organ-level immune alterations preceding clinical onset.

### IN-based Stratification Enables Identification of Macrophage Associated TNF-**α**/NF-**κ**B Program Related to T1D Pathogenesis

The IN based T1D gene signature was subsequently leveraged to stratify NOD mice at the early and intermediate stages and investigate the distinct pancreas transcriptomic profiles of progressors and non-progressors. Single cell sequencing performed on the pancreas at early and intermediate stage was analyzed for corresponding transcriptomic differences in immune cells (**Figure 4a)**. Mice classified as progressors or non-progressors by the SVC model were selected for downstream pancreatic analyses (**see Methods, Supplementary Figure 7**).

**Figure 4.**
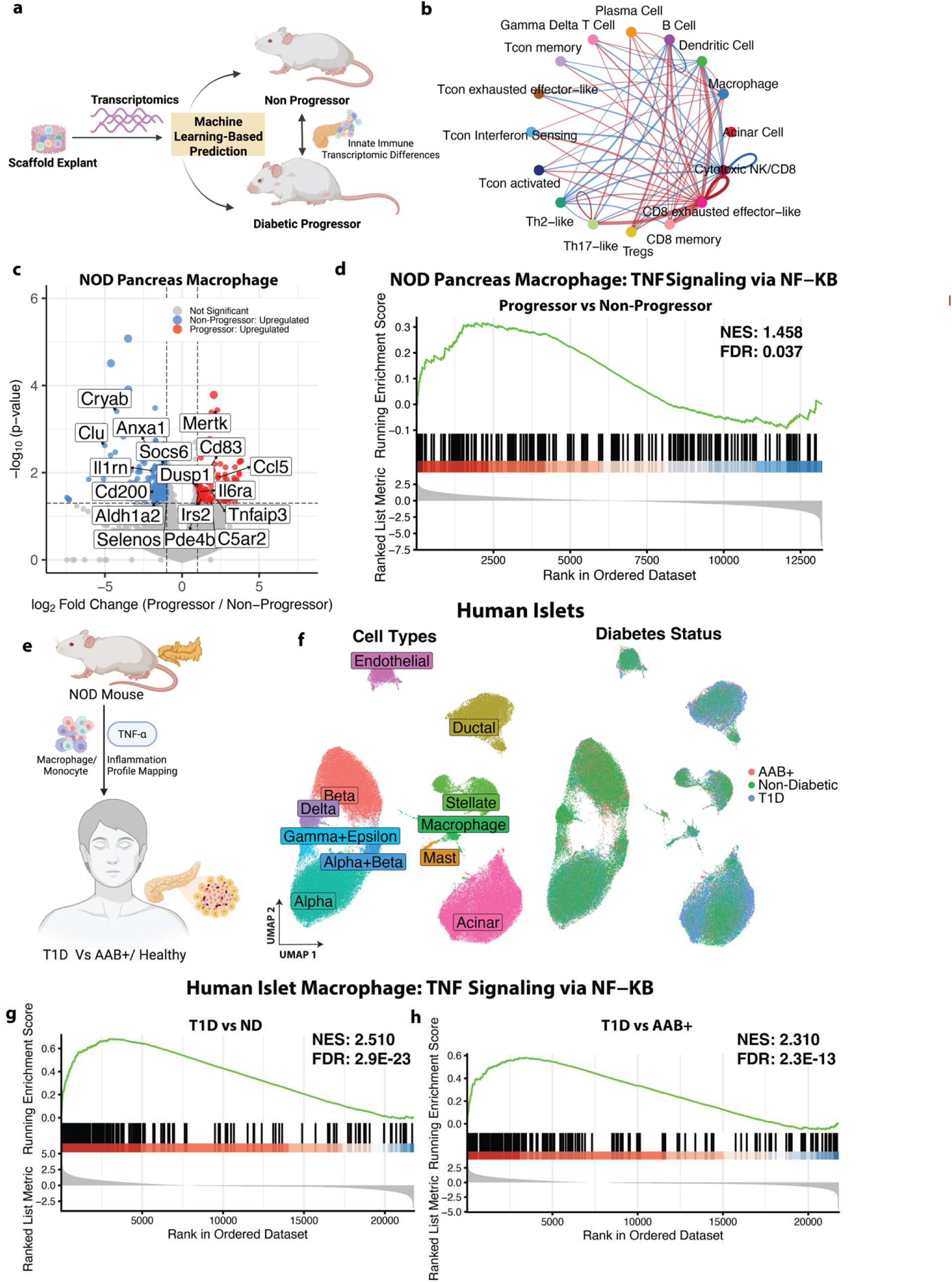
IN-based stratification reveals TNF-α related pro-inflammatory macrophage program related to T1D progression in mouse and humans. **a,** IN-based T1D gene signature were used to stratify progressor (n=6) and non-progressor(n=3) NOD mice from early and intermediate stages to investigate differences in immune transcriptomics in the pancreas **b,** CellChat analysis of the pancreatic microenvironment showing top 25% increased (red) and decreased (blue) cellular communication probabilities in progressors versus non-progressors, highlighting enhanced macrophage-associated interactions. **c,** Volcano Plot showing pancreas macrophage DEGs (p ≤ 0.05, |log₂FC| ≥ 1) between progressors vs non-progressors NODs. Axes were constrained for visualization (log₂FC: −8 to 8; –log₁₀(p): 0 to 6) **d,** GSEA Enrichment plot of TNF-*α* pathway in pancreas macrophage in the NOD progressor vs non-progressors. Normalized Enrichment Score (NES): 1.458, FDR q-value:L0.037. **e**, TNF-α-associated program identified in pancreas macrophage in NOD progressors was examined in human islet macrophages to determine whether a similar inflammatory signature is present in T1D patients compared with autoantibody positive (AAB+) and non-diabetic (ND) individuals. **f,** UMAP of human pancreas islets grouped by cell types and patient status-T1D (n=10), AAB+ (n=11), ND (n=27). **g-h,** GSEA Enrichment plot of TNF*α* pathway in pancreas macrophage in **g**, T1D vs ND individuals. (NES: 2.510, FDR q-value:L2.9E-23) and **h**, T1D vs AAB+ individuals (NES: 2.310, FDR q-value:L2.3E-13).

Alterations in intercellular communication within the pancreatic immune microenvironment between predicted progressors vs non-progressors were analyzed using CellChat analysis (**Figure 4b**). CD8 exhausted effector-like T cells exhibited higher interaction probabilities in the progressors compared to non-progressors, consistent with the role of exhausted cytotoxicity-related CD8 T cells in T1D reported by prior works (26). Notably, among innate immune populations, macrophages demonstrated increased interaction probability in progressors. This increased interaction of macrophages within the pancreas immune microenvironment and their high representation in the early-stage IN immune microenvironment suggested a key role for macrophage-mediated signaling in shaping T1D disease progression. Accordingly, for the subsequent analysis, we primarily focused on transcriptional differences in pancreatic macrophages to identify inflammatory programs underlying their enhanced interaction in progressors vs non-progressors.

Differential expression analysis identified 362 DEGs, highlighting substantial transcriptomic divergence in macrophages from progressors versus non-progressors beginning at early and intermediate disease stages (**Figure 4c, Supplementary Table 7)**. Interestingly, many TNF-α signaling related inflammatory cytokine genes like *Il6ra*, *Tnfaip3*, *Cd83*, *Ccl5*, *Pde4b*, *Dusp1*, and *Mertk* were upregulated in progressor macrophages, suggesting heightened inflammatory activation **(Figure 4c)**. In contrast, genes associated with immune regulation like *Clu*, *Cryab*, and *Cd200* were downregulated. Collectively, these DEGs suggested that macrophages in T1D progressors exhibit heightened TNF-α related signaling. This observation was further confirmed by GSEA which identified TNF-α Signaling via NF-κB signaling as a significantly enriched pathway in T1D progressors vs non-progressors **(Figure 4d)**. These findings suggest that macrophages may contribute to heightened inflammation in the pancreatic microenvironment through TNF-α/NF-κB driven pathway in T1D progressors vs non-progressors. We assessed the conservation of the macrophage-associated TNF-α program identified in NOD progressors in human T1D. We investigated the transcriptomics profile of islet resident macrophages from individuals with T1D, autoantibody positive (AAB+) donors and non-diabetic (ND) controls (**Figure 4e**). Across all three groups, macrophage populations were identified within human islets, along with other endocrine cells (**Figure 4f**). GSEA revealed that TNF-α signaling via NF-κB was significantly upregulated in macrophages from T1D individuals compared to both ND and AAB+ controls (**Figure 4g, h**). The persistence of this TNF-α enrichment in T1D versus AAB+ macrophages indicated that TNF-α related inflammation is not merely a feature of autoimmunity or autoantibody positivity but is specifically associated with progression to overt diabetes.

### IN-Based response score can identify resistance to anti-TNF***α*** treatment

The prominent role of TNF-α in the T1D pathogenesis was next evaluated through pharmacologic blockade of TNF-α and subsequent determination of disease trajectory. NOD mice received anti-TNF-α treatment (100 μg i.p., every other day, 10 doses) initiated at an early (Week 6) or intermediate (Week 12) stage, with untreated mice serving as controls. INs were explanted at pre- and post-treatment timepoints to monitor the transcriptomic profile (**Figure 5a**). Anti-TNF-α therapy conferred partial protection from T1D onset at both treatment stages.

**Figure 5:**
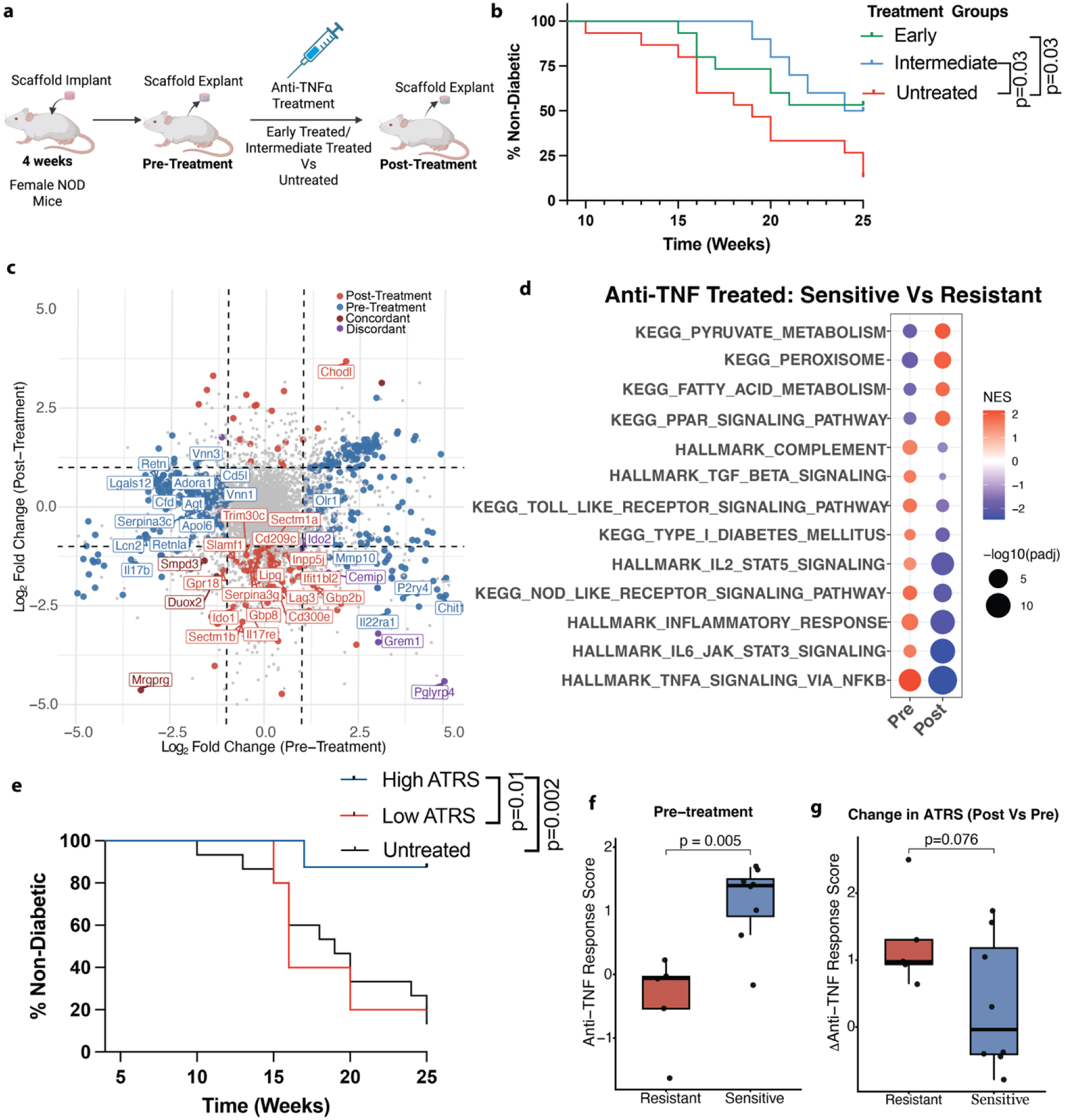
IN Based Response Score Can Identify Resistance to Anti-TNF-*α* Treatment. **a,** Experimental Design: Mice were treated with anti-TNF-α at early (n=15) and intermediate (n=10) stage and compared to untreated control (n=15). IN were explanted at pre-treatment and post-treatment timepoint. **b,** Survival curve shows percentage of non-diabetic in the anti-TNF-α treated early, intermediate and untreated control groups till 25 weeks of ages. Statistical analysis was done using log-rank Mantel-Cox test. **c,** Double volcano plot depicting differentially expressed genes within the IN between Sensitive (n=8) and Resistant (n=5) mice which were given early anti-TNF-α treatment (p ≤ 0.05, |log₂FC| ≥ 1). The x-axis represents log₂FC of Sensitive vs Resistant at the pre-treatment timepoint, and the y-axis represents log₂FC at the post-treatment timepoint. Axes were constrained to ±5 log₂FC for visualization. **d,** Gene Set Enrichment Analysis showing selected upregulated and downregulated pathways in the early anti-TNF-α treated sensitive vs resistant group at pre-treatment and post-treatment. **e,** T1D Incidence in anti-TNF treated mice with High (n=8) and Low (n=5) Anti-TNF Response Score (ATRS) at pre-treatment vs untreated. Statistical analysis was done using log-rank Mantel-Cox test **f,** ATRS in sensitive vs resistant mice at pre-treatment timepoint **g,** Change in ATRS in sensitive vs resistant mice from pre- to post-treatment timepoint. Statistical analysis was performed using Welch’s t-test.

By 25 weeks of age, ∼54% (8/15) of early-treated and 50% (5/10) of intermediate-treated mice remained normoglycemic, compared to only 13% (2/15) in the untreated group (**Figure 5b, Supplementary Table 8**). These results indicated that while TNF-α blockade can delay or prevent T1D progression, there is variation in response.

Given the heterogeneous efficacy of anti-TNF-α therapy, we identified an IN related transcriptional score to stratify resistant from sensitive groups. We compared niche transcriptomes before and after anti-TNF-α treatment in the early-treatment cohort (**Supplementary Table 9**). Anti-TNF-α sensitive mice exhibited a distinct transcriptional profile compared to the resistant animals. Inflammatory genes like *Grem1* and *Cemip*, were elevated at pre-treatment in sensitive mice and reduced after therapy, indicating reversal of an inflammatory program in sensitive animals (**Figure 5c**). Consistently, post anti-TNF-α treatment, sensitive mice demonstrated downregulation of multiple immune activation genes, including *Il17re*, *Gbp8*, *Cd209c*, *Slamf1*, and *Cd300e*, reflecting attenuation of inflammatory signaling following TNF-α blockade. (**Figure 5c**). At the pathway-level, pre-treatment, sensitive mice showed enrichment of immune and inflammation related pathways, including inflammatory response, TNF-α via NFK-*β*, IL-6 JAK STAT3, IL2-STAT5, Toll and NOD-like receptor signaling, TGF-β signaling and complement pathway, consistent with a state of heightened cytokine-driven inflammation (**Figure 5d, Supplementary Table 9**). Following anti-TNF-α therapy, these inflammatory pathways exhibited relative attenuation in sensitive mice compared with resistant animals. In contrast, peroxisome-associated pathways were diminished pre-treatment but relatively enriched following therapy in sensitive mice (**Figure 5d**). Collectively, these findings indicated that underlying immune-inflammatory heterogeneity at baseline dictated the efficacy of anti-TNF-α therapy.

Building on these pathway-level differences, we next investigated the pathway level programs to distinguish sensitive and resistant mice by developing a quantitative metric to predict therapeutic response. Using Gene Set Variation Analysis (GSVA) scores derived from pre-treatment IN transcriptomes, we applied an elastic net model to identify a minimal set of pathways that best discriminated anti-TNF sensitive from resistant animals. This analysis yielded an Anti-TNF-α Response Score (ATRS), integrating weighted GSVA scores from selected inflammation related pathways (HALLMARK TNF-α signaling via NF-κB, HALLMARK TGF-β signaling, KEGG Peroxisome). When the mice were stratified by baseline ATRS, 87.5% (7/8) of animals with a high baseline ATRS remained normoglycemic till 25 weeks of age (**Figure 5e**). In contrast, only 20% (1/5) of those mice with low baseline ATRS remained normoglycemic over the same period (**Figure 5e**). Consistent with this observation, resistant mice exhibited significantly lower ATRS pre-treatment compared to the sensitive ones (**Figure 5f**). Following treatment, the sensitive mice demonstrated a trend toward greater reduction of ATRS from pre- to post-treatment than the resistant ones, reflecting modulation of inflammatory state in sensitive mice (**Figure 5g**). Together, these results indicate that the IN derived ATRS captures baseline immune inflammatory differences and treatment associated changes between sensitive and resistant mice, highlighting the diagnostic ability of the IN for identifying therapeutic responsiveness to anti-TNF-α.

## Discussion

Early immune dysregulation within disease-relevant tissues is a key feature of T1D pathogenesis and a critical determinant of immunotherapy efficacy (27, 28). Yet current clinical monitoring strategies rely largely on glucose measurements and circulating autoantibodies, which reflect downstream metabolic outcomes and incompletely capture the upstream inflammatory programs. Increasing evidence also suggests that tissue-resident immune dynamics differ substantially from circulating blood profiles, limiting the ability of peripheral markers to resolve early pathogenic inflammation (29–31). To address this gap, we leveraged an implantable biomaterial scaffold as a cell capture platform, that vascularizes to form a synthetic immunological niche, enabling minimally invasive access to tissue-relevant immune activity related to T1D pathogenesis.

The IN captured dynamic immune remodeling in T1D, revealing a temporal shift from early myeloid-related dysregulation, dominated by macrophages, to progressively increased T cell-associated perturbations as disease progressed. Comparisons of progressors and non-progressors within T1D-associated vital tissues at early preclinical stages have been limited, largely due to the inability to prospectively distinguish divergent disease trajectories. The IN overcomes this barrier by enabling longitudinal characterization of tissue-specific immune differences between progressors and non-progressors across T1D stages. At early stages, the IN had a higher proportion of macrophages followed by increased T cell representation at later stages. Consistent with the flow results, the early stage IN-based T1D gene signature reflected a predominantly innate inflammatory state with integrated adaptive elements. Prominent innate components included macrophages and monocyte associated genes like *Cpvl and Tm9sf4* implicated in antigen presentation and inflammation, *Psme3*, a NF-κB inflammation-associated gene and complement pathway genes like *Cfi,* involved in islet autoimmunity (32–36). In addition, the signature also had adaptive immune regulator genes like *Cd40lg* which has previously been reported as therapeutic target for reducing T1D inflammation, *Ccr8,* a gene reported to be a master drivers of immune Treg immune modulation, and *Pdcd1,* a key signature of immune exhaustion related to T1D pathogenesis and treatment efficacy (37–41). As T1D progressed to the intermediate stage, the IN transcriptomic landscape showed increased representation of adaptive immune DEGs, including interferon-responsive and cytotoxic genes, along with enrichment of T cell-associated pathways such as IFN-α/γ signaling. This increase in adaptive dysregulation was accompanied by enrichment in broader inflammatory networks like TNF and IL-6 signaling in progressors vs non-progressors. Together, the findings from the IN support a staged model in which early myeloid-skewed inflammation establishes a permissive immune niche that later integrates adaptive immune programs related to interferon and cytotoxicity (42, 43).

The IN-derived T1D gene signature captured tissue immune pathology and distinguished T1D from controls in disease-relevant tissues, but not in peripheral blood. Many genes within the IN signature mapped to key innate and adaptive immune populations in the NOD pancreas and corresponding human T1D tissues, including spleen and pancreatic lymph node whereas a substantial proportion were not detected in human PBMCs. Accordingly, human orthologs of this signature separated T1D from non-diabetic samples in spleen and pancreatic lymph node but demonstrated limited discriminatory capacity in blood. This tissue-associated performance indicates that the IN-derived T1D gene signature reflects immune-inflammatory processes active within disease-relevant tissues that are incompletely represented in circulation. These tissue-specific T1D changes are consistent with previous studies that have reported T1D-related immune cell type differences present in spleen and lymph nodes but absent in peripheral blood, highlighting compartmentalized immune responses in T1D (44–47).

The early-stage IN-derived T1D gene signature stratified progressors from non-progressors and identified pancreatic macrophages as a key population marked by heightened TNF-α/NF-κB signaling in progressors. TNF-α-mediated inflammation in pancreas macrophages has previously been reported as a key feature of T1D pathogenesis with TNF-α promoting antigen presentation and potentiating insulitis (48–50). Genetically disrupting TNF signaling has shown to protect NOD mice from the onset of diabetes and reduce insulitis, supporting a causal role for TNF pathways in disease progression (51). The TNF-α signature was further conserved in human T1D, where islet macrophages from T1D individuals showed significantly elevated TNF-α inflammation relative to those from autoantibody-positive and non-diabetic individuals, indicating that TNF-α amplification is specifically linked to progression toward overt clinical disease rather than autoimmunity alone. Consistent with its pathogenic role in T1D, therapeutic TNF-α blockade has preserved residual β-cell function in subsets of patients, as demonstrated in new-onset trials with etanercept and golimumab, supporting TNF-α as a clinically actionable yet heterogeneously effective target (52, 53).

Analysis of the IN identified key T1D mediators that, upon therapeutic targeting of TNF-α, altered the disease trajectory and enabled pre-treatment stratification of resistant and sensitive individuals. Anti-TNF-α treatment has been reported to partially reduce diabetes incidence in NOD mice, though significant heterogeneity in the therapeutic response was observed, consistent with prior studies (54, 55). Leveraging IN-derived transcriptomic profiles, we identified the immune and inflammatory programs underlying this variability in response. Anti-TNF-α was more effective in sensitive mice with heightened TNF-α inflammation at baseline, accompanied by upregulated TGF-β signaling and downregulated peroxisome related pathways as compared to the resistant ones. While TGF-β signaling has been implicated in inflammatory regulation and autoimmunity in T1D, peroxisome-related pathways linked to PPAR-γ activity have been associated with β-cell protection and metabolic resilience in NOD models (56, 57). Building on these dynamics, we developed a pathway-integrated Anti-TNF-α Response Score (ATRS) that quantitatively captures the changes in TNF-α, TGF-β, and peroxisome-associated signaling programs within the IN. ATRS discriminated between sensitive and resistant mice at baseline and enabled pre-treatment stratification of therapeutic responsiveness. These findings suggest that IN-derived ATRS may provide a framework for stratifying candidates most likely to benefit from TNF-α-targeted intervention— an important consideration given the variable efficacy of anti-TNF-α therapies like golimumab, in T1D clinical trials (54). The heterogeneity in inflammatory and immune states captured by the IN may also facilitate the decision for which therapeutic to administer. Early-stage inflammation-dominant profiles may respond sufficiently to TNF-α blockade alone, whereas individuals with more persistent autoreactive immune programs in later stages may be more resistant to anti-TNF-α and require combinatorial approaches that integrate anti-inflammatory therapy with T cell-directed agents such as anti-CD3. In this context, the IN functions as a precision medicine platform capable of tailoring immunomodulatory strategies to individual immune states.

In conclusion, we demonstrate that the IN captures underlying stage wise immune inflammatory programs that distinguish T1D progressors from non-progressors in NOD mice and are conserved in human T1D. Mice progressing to diabetes exhibited early innate immune dysregulation, whereas the adaptive responses increased as disease progressed. At early stages, macrophages were enriched in a TNF-α inflammatory profile in T1D progressors. Anti-TNF-α delayed disease onset, yet heterogeneity in therapeutic response was observed. This heterogeneity was captured by the IN and quantified through a pathway response score that distinguished treatment-sensitive from resistant individuals. Collectively, these findings establish the synthetic IN as a minimally invasive platform for detecting stage-specific immune-inflammatory perturbations and monitoring therapeutic response, with potential utility across immune-mediated disorders.

## Methods

### Sex as a biological variable

Female non-obese diabetic (NOD/ShiLtJ, strain #001976; The Jackson Laboratory) mice were used due to their higher incidence of type 1 diabetes (58). Prior studies indicate that the underlying immune mechanisms are conserved across sexes (59). Both sexes were included for human tissues.

### Study approval

All animal studies were conducted in accordance with institutional guidelines and protocols (PRO00011484, PRO00011621) approved by the University of Michigan Institutional Animal Care and Use Committee.

### Animal Cohort Design

For T1D monitoring cohorts, the immunological niche was implanted subcutaneously at 4 weeks of age. Niches were explanted and replaced at defined disease stages-early (6 weeks), intermediate (12 weeks), and late (18 weeks) for downstream processing and analysis. For the pancreas single-cell profiling study, niches were implanted at 4 weeks of age and explanted at 6 weeks of age to generate transcriptomic profile for disease progression prediction. Pancreata from the mice were harvested at the early (6 weeks) and intermediate (12 weeks) time points for immune cell isolation and single-cell RNA sequencing.

For the EAE model, the INs were used in murine experimental autoimmune encephalomyelitis as previously described (60). In brief, EAE was induced by adoptive transfer of T cells reactive to myelin oligodendrocyte glycoprotein (MOG) peptide 35 to 55 and control animals received an adoptive transfer of T cells reactive to ovalbumin (OVA) peptide 323 to 339. Mice received a subcutaneously implanted IN 14 days before the adoptive transfer of T cells on Day 0. Scaffolds were explanted 6 days after the adoptive transfer before clinical symptoms were present. For the 4T1 model, female BALB/c mice (7-8 weeks) underwent subcutaneous scaffold implantation followed by orthotopic mammary fat pad inoculation 4T1 cells, with tumor progression monitored for 21 days, as previously described (21). The adoptive transfer T1D model was established by activating splenocytes from 6-week-old NOD BDC2.5 donor mice with cognate peptide in vitro, followed by intraperitoneal transfer of 5×10L activated cells into age-matched NOD.scid recipients, as previously described (20). The INs were explanted at the time of transfer and 10 days post-transfer.

For anti-TNF-α treatment studies, mice were treated beginning at either 6 weeks of age (early treatment) or 12 weeks of age (intermediate treatment). Animals received intraperitoneal injections of 100 μg anti-TNF-α antibody (clone XT3.11; Bio X Cell, #BE0058) every other day for a total of 10 doses. Immunological niches were explanted before treatment initiation (6 weeks) and after treatment completion (9 weeks for early-treated and 15 weeks for intermediate-treated groups) for downstream molecular profiling.

### Glucose Monitoring and Intraperitoneal Glucose Tolerance Test

Non-fasted blood glucose was measured weekly by tail-vein sampling using an Accu-Chek Aviva Plus glucometer and test strips (Roche, Catalog No. 6908373001). Diabetes incidence was defined as two consecutive blood glucose readings ≥250 mg/dL or study completion at 30 weeks of age. For intraperitoneal glucose tolerance testing (IPGTT), mice were fasted for 4 h, then injected intraperitoneally with sterile 20% (w/v) D-glucose in PBS (2 g/kg body weight).

Blood glucose was measured from the tail vein at 0 (pre-injection), 15, 30, 45, 60, and 120 min post-injection. Glucose tolerance was quantified from the glucose excursion curve and by area under the curve (AUC). The blood glucose and IPGTT values are in **Supplementary Table 1**.

### Immunological Niche fabrication

Microporous poly(ε-caprolactone) (PCL) based immunological niches were fabricated by salt leaching technique as previously described (13, 17, 18). First 99g of NaCl (250 to 425um) was mixed with 1g of PCL pellets for 1hour at 85 °C. Subsequently, around 77.5 mgLof the PCL-NaCl melt dispersion was pressed into 5mm scaffolds at 1500Lpsi for 45 seconds. They were then heated at 65<C for 5 mins on each side for annealing of PCL. The scaffolds were placed in deionized water bath for salt leaching, and the resulting porous niches were sanitized with 70% ethanol immersion followed by sterile PBS rinses.

### Immunological Niche implantation and retrieval

Mice were anesthetized with 2% isoflurane and maintained under anesthesia via a nose cone throughout all surgical procedures. The immunological niche was implanted into the subcutaneous space through a small dorsal incision. To minimize post-operative pain, mice received carprofen (5 mg/kg, subcutaneous) at the time of surgery and again 24Lhours post-operatively and were monitored during recovery for signs of distress. For the IN retrieval, mice were re-anesthetized with 2% isoflurane, and the implant site was shaved and disinfected. A small incision adjacent to the implant was reopened using sterile forceps, and the IN was carefully excised and removed. The incision was closed with wound clips.

Explanted niche designated for transcriptomics, metabolomics or lipidomics analysis were immediately flash-frozen in isopentane, maintained on dry ice, and stored at −80 °C until further processing. Niches intended for flow cytometry were immediately placed in RPMI 1640 medium (Gibco) and stored in ice prior to processing.

### Flow Cytometry

The explanted immunological niches were prepared for flow cytometry by mechanical dissection and enzymatic incubation. Briefly, the niches were minced and then digested in Liberase,TL (Millipore Sigma, Catalog No.5401020001) at 10 U/mL for 30 minutes at 37°C. Following digestion, the niches were mashed through a 70-µm strainer, lysed for red blood cells using ACK buffer (Fisher, Catalog No. A1049201) for 1 minute and then the remaining cells extensively washed with fluorescence-activated cell sorting buffer: PBS with 0.5% bovine serum albumin (Sigma-Aldrich, St. Louis, MO) and 2 mM EDTA. IN from each mouse was equally split into two parts to enable staining and analysis of myeloid and lymphoid immune cells from the same sample. They were then blocked with anti-CD16/32 (1:50, Biolegend, Catalog No. 156604). The cells were stained with the myeloid antibodies and lymphocyte antibodies for their respective panels using dilution mentioned in Table 1 and 2 respectively. Flow cytometry was performed using a Cytek Northern Lights Aurora. Flow data was analyzed with FlowJo v10.8.2. Single stain and FMO controls were run with each timepoint condition to aid gating and compensation (**Supplementary Figure S1**).

**Table 1:** Myeloid Panel Dyes.

| Dye | Marker | Catalog Number | Dilution | Vendor |
| --- | --- | --- | --- | --- |
| Alexa Fluor 700 | CD45 | 103128 | 1:100 | BioLegend |
| Brilliant Violet 510 | CD11b | 101245 | 1:10 | BioLegend |
| Brilliant Violet 785 | CD11c | 117335 | 1:40 | BioLegend |
| FITC | Ly6C | 128006 | 1:100 | BioLegend |
| Brilliant Violet 711 | F4/80 | 123147 | 1:20 | BioLegend |
| APC | CD86 | 159216 | 1:80 | BioLegend |
| Brilliant Violet 605 | CD206 | 141721 | 1:10 | BioLegend |
| PE | CD170 | 164104 | 1:40 | BioLegend |
| PE/Cyanine 7 | CD163 | 155320 | 1:40 | BioLegend |
| Brilliant Violet 650 | CD274 | 124336 | 1:20 | BioLegend |
| Zombie Violet | Viability | 423114 | 1:50 | BioLegend |

**Table 2:** Lymphoid Panel Dyes.

| **Dye** | **Marker** | **Catalog Number** | **Dilution** | **Vendor** |
| --- | --- | --- | --- | --- |
| Alexa Fluor 700 | CD45 | 103128 | 1:100 | BioLegend |
| Brilliant Violet 650 | CD3 | 100229 | 1:10 | BioLegend |
| PE | CD4 | 100408 | 1:70 | BioLegend |
| Spark Blue 574 | CD8 | 100794 | 1:100 | BioLegend |
| Brilliant Violet 510 | CD279 (PD1) | 135241 | 1:40 | BioLegend |
| Brilliant Violet 711 | CD223 (LAG3) | 125243 | 1:20 | BioLegend |
| PE/Cyanine 7 | CD25 | 101915 | 1:30 | BioLegend |
| Zombie Violet | Viability | 423114 | 1:50 | BioLegend |

### Bulk RNA Sequencing Analysis

Frozen immunological niche were lysed in 800 µL of TRIzol reagent (Fisher Scientific, Cat. No. 15596026) and homogenized at 15,000 rpm. RNA was isolated from the homogenized solution with a direct-zol RNA Miniprep Plus (Fisher Scientific, Cat. No. NC1047980) kit following the manufacturer’s instructions. The concentration of the isolated RNA was measured using a NanoDrop and the samples were submitted to the Advanced Genomics Core at the University of Michigan for bulk RNA sequencing. Library preparation was performed using ribosomal RNA depletion for total RNA library construction. Sequencing was carried out on the Illumina NovaSeqXPlus. BCL Convert Conversion Software v4.0 (Illumina) was used to generate de-multiplexed Fastq files.

Raw gene-level count matrices from three independent bulk RNA-seq cohorts were jointly processed for differential gene expression (DEG) analysis. Data for two of these cohorts were obtained from previously published work in King et al. Samples were stratified by disease stage as follows: early (6 weeks of age; 16 non-progressors and 24 Progressors), intermediate (12-14 weeks; 10 non-progressors and 10 progressors), and late (17-20 weeks; 4 non-progressors and 10 progressors). All downstream analyses were performed independently within each disease stage. For each stage, genes that were unexpressed across all samples or sparsely detected (zero counts in >85% of samples) were removed. To restrict analyses to well-annotated and biologically interpretable transcripts, genes annotated as RIKEN or pseudogenes were excluded prior to differential expression testing. Differential expression analysis was performed using DESeq2 (v1.50.2), incorporating cohort batch as a covariate in the design formula. Log₂ fold changes were estimated for progressors relative to non-progressors using the Wald test. Genes with an absolute log₂ fold change ≥ 1 and Wald p-value ≤ 0.05 were considered differentially expressed. Gene set enrichment analysis (GSEA) was performed separately for each disease stage using clusterProfiler, with gene sets obtained from MSigDB, including Hallmark and KEGG pathway collections (61). All the genes were ranked by the DESeq2 Wald statistic (Progressor vs. Non-Progressor) and pathways with adjusted p ≤ 0.1 and absolute normalized enrichment score (|NES|) ≥ 1 were considered significantly enriched. Complete differential expression and GSEA results for all stages are provided in **Supplementary Table 3**. For visualization of pathways, a curated subset of immune, signaling, and metabolic pathways that were significantly enriched in at least one disease stage were selected.

To assess the disease specificity of differentially expressed genes (DEGs), we analyzed niche-associated transcriptomic datasets from additional inflammatory disease models and compared them to the early stage T1D niche signature. For the 4T1 breast cancer model, bulk RNA-seq data were obtained from Orbach *et al.* and differential expression at niche was assessed between healthy and tumor-bearing mice at the pre-symptomatic Day 14 timepoint (21). For the Experimental Autoimmune Encephalomyelitis (EAE) model, niche samples collected at the pre-symptomatic Day 6 stage were similarly compared between diseased and healthy conditions.

DEGs and enriched pathways for these models were identified using the same analytical pipeline described above. DEGs and pathway enrichment results from these datasets were compared specifically to the early-stage T1D DEGs. In addition, for comparison with a highly T cell-driven T1D context, adoptive transfer model data were obtained from King *et al.*, in which NOD.scid mice received either diabetogenic BDC2.5 T cells or ovalbumin (OVA) control T cells. Differential expression analysis was performed on niche samples collected at day 10 post-transfer, and the resulting gene and pathway signatures were compared against early-stage T1D results. DEGs and enriched pathways from the 4T1 cancer, EAE and Adoptive Transfer T1D model are available in **Supplementary Table 4**.

For identifying an early transcriptional signature predictive of T1D progression, filtered and limma based batch corrected genes that were significantly differentially expressed between progressor and non-progressor at the early stage (Welch p-value ≤ 0.05) were selected as input features to a Partial Least Squares–Discriminant Analysis (PLS-DA) model (62). Top 100 genes with the highest Variable Importance in Projection (VIP) scores were selected as the core discriminatory signature, which robustly separated progressors from non-progressors at the early timepoint.

This gene signature was subsequently used to train a Support Vector Classifier (SVC) model to predict T1D progression at early stage. All machine learning analyses were conducted in Python (v3.11.1) using scikit-learn (v1.5.1) and pandas (v2.2.2). The dataset comprising 40 samples was stratified into a training and validation set into a 7:3 ratio using a stratified sampling method to preserve class balance. Model hyperparameters were optimized using a 5-fold cross validation on the training data and a linear SVC kernel, with regularization parameter C = 1 was selected. Model performance was subsequently evaluated across 1000 independent stratified train validation sets to assess robustness. This model was used to infer progression status of animals used for the pancreas single-cell sequencing using their early-stage niche-derived bulk RNA-sequencing profile. Out of 10 mice (6 harvested for pancreas was extracted at early timepoint and 4 for intermediate timepoint), 9 were confidently (>60% probability) predicted as progressor or non-progressor and used for downstream analysis of single cell RNA analysis. The progression assignments are summarized in **Supplementary Figure 7.**

Differential gene expression and gene set enrichment analyses for the early anti-TNF-α-treated cohort were performed using the same pipeline described above. To derive an Anti-TNF-α Response Score (ATRS), pathway-level activity scores were computed on a per-sample basis using Gene Set Variation Analysis (GSVA). GSVA scores for pathways associated with inflammatory signaling and immune-metabolic regulation that were significantly enriched (adjusted p ≤ 0.1 and |NES| ≥ 1) in both pre- and post-treatment comparisons were used as input features for an elastic-net logistic regression model (α = 0.7), trained to distinguish anti-TNF responders from non-responders at the early treatment time point. Model regularization and feature selection were performed using leave-one-out cross-validation. The final model selected three pathways-KEGG_PEROXISOME, HALLMARK_TGF_BETA_SIGNALING, and HALLMARK_TNFA_SIGNALING_VIA_NFKB-which together defined the ATRS. The resulting model assigned positive weights to TNF-α/NF-κB and TGF-β signaling and a negative weight to peroxisome metabolism. Sample-level ATRS values were calculated as a linear combination of GSVA pathway scores weighted by the corresponding model coefficients. Full input pathway selection list, model coefficients, and ATRS values are provided in **Supplementary Table 10**.

### Single Cell RNA Sequencing

The explanted pancreatic tissues were mechanically dissected and digested in collagenase for 15 minutes at 37°C. Following digestion, the samples were mashed through a 70-µm strainer, lysed for red cells using ACK buffer (Fisher, Catalog No. A1049201) and washed with fluorescence-activated cell sorting buffer: PBS with 0.5% bovine serum albumin (Sigma-Aldrich, St. Louis, MO) and 2 mM EDTA to dissociate into single cell suspensions for subsequent immune cell enrichment. Cells were stained with AF700-CD45 (BioLegend) and CD45+ populations were isolated using flow-assisted cell sorting on the Bigfoot Cell Sorter (Invitrogen) (**Supplementary Figure 5**). Following sorting, cells were fixed according to the manufacturer’s protocol for the 10× Chromium Next GEM Single Cell Fixed RNA Sample Preparation Kit. Fixed samples were submitted to the University of Michigan Advanced Genomics Core, where libraries were constructed using the 10× Single Cell Fixed RNA Hybridization & Library Kit in combination with the Single Cell Fixed RNA Mouse Transcriptome Probe Kit, following manufacturer guidelines for the 10× Single Cell Gene Expression Flex platform. Final Samples were sequenced on the NovaSeq X 10B flow cells (300 cycle) with a target depth of approximately 25,000 reads per cell.

For each sample, filtered and unfiltered CellRanger (v8.0.0) outputs were processed independently prior to downstream integration. Ambient RNA contamination was mitigated using SoupX, followed by quality control filtering with ddqcR and removal of putative doublets using DoubletFinder (63, 64). Processed datasets were subsequently merged, normalized using SCTransform, and integrated according to the standard workflow implemented in Seurat v5 (65, 66). The preprocessing and analysis pipeline is publicly available on GitHub.

Dimensionality reduction was performed using principal component analysis (PCA), retaining 30 principal components. Cells were embedded in two dimensions using uniform manifold approximation and projection (UMAP) based on the same number of dimensions and clustered. Cell clusters were annotated using previously established marker genes (**Supplementary Figure 5**) (25, 67). For downstream analysis, only mice which were predicted with >60% confidence by the SVC model as T1D progressors or non-progressors were included (**Supplementary Figure 7**). This yielded five mice at the early time point (4 Progressor, 1 Non-Progressor) and four mice at the intermediate time point (2 Progressor, 2 Non-Progressor). Downstream analysis was performed for progressor vs non-progressors at both the timepoints combined.

Intercellular communication was inferred using CellChat to model ligand-receptor signaling differences between T1D progressors and non-progressors at early and intermediate stages combined (68). For the macrophage differential gene expression analysis, the macrophage data was pseudo bulked by summing across macrophage cells within each mouse and DESeq was conducted at the sample level treating time as a co-variate. Gene Set enrichment analysis was performed as previously described.

### Human Validation

The human PBMC single cell RNA sequencing data was obtained from Hoonardoost *et al.* and annotated using previously identified markers (69). The pancreatic islet dataset was obtained from PanKbase and the previously identified annotations were used for the downstream analysis (70).

HPAP spleen and PLN snMultiome (RNA+ATAC) datasets were analyzed using RNA-based downstream workflows, with joint RNA–ATAC metrics informing nucleus-level, donor-adaptive quality control (71). Detailed alignment, decontamination, multimodal QC, integration, and annotation procedures are described in the Supplementary Methods.

For the PLS-DA clustering using human orthologs of T1D gene signature, the single cell datasets were pseudo bulked by summing gene counts across cells within each sample, followed by normalization to the mean counts per cell. For GSEA enrichment of macrophage related TNF in islet in NOD and human islets, the data was pseudo bulked by summing counts across monocyte cells within each donor group sample.

### Metabolomics and Lipidomics Analysis

Immunological niche samples were bead-homogenized and extracted in 80% methanol for targeted metabolomics. Supernatants were weight-normalized, dried under vacuum, and reconstituted in 50% methanol prior to analysis. Metabolites were quantified using an Agilent 1290 Infinity II LC coupled to a 6495d QqQ mass spectrometer with dynamic multiple reaction monitoring (dMRM). Separation was performed on a Poroshell 120 HILIC-Z column. The method was optimized to detect 435 metabolites in both positive and negative ion modes. Detailed solvent compositions and gradient parameters are provided in the Supplementary Methods.

For lipidomics, samples were extracted using a chilled biphasic MTBE:methanol system. The organic phase was analyzed on an Agilent 1290 Infinity II LC–6495d QqQ platform using a ZORBAX Eclipse Plus C18 column with a targeted dMRM acquisition method optimized for 665 lipid species in positive and negative ion modes. Chromatographic gradients and solvent compositions are described in the Supplementary Methods.

Raw data were processed using Agilent MassHunter Quantitative Analysis (v12.1). Data were median-centered and pareto-scaled, followed by fold-change analysis and Student’s *t*-tests relative to control groups using MetaboAnalyst (v5.0) (72).

### Statistics

Plots for flow data, IPGTT, blood glucose and survival were generated, and statistical analyses were performed using GraphPad Prism 10.6.1 (GraphPad Software). Survival curves were created by Kaplan-Meier survival analysis combined with the log-rank Mantel-Cox test.

Statistical analysis for flow data was performed using a mixed-effects model with the Geisser–Greenhouse correction, followed by Tukey’s multiple-comparison test, with individual variances computed for each comparison. For IPGTT, statistical analysis was performed using multiple unpaired t-tests with single pooled variance.

For bulk and pseudo-bulk RNA sequencing of the IN and pancreas macrophages, differential expression analysis was performed using DESeq2 (v1.50.2). Log₂ fold changes were estimated for progressors relative to non-progressors using the Wald test. Genes with an absolute log₂ fold change ≥ 1 and Wald p-value ≤ 0.05 were considered differentially expressed. For GSEA analysis, clusterprofiler was used and pathways with adjusted p ≤ 0.1 and absolute normalized enrichment score (|NES|) ≥ 1 were considered significantly enriched. For Anti-TNF response score comparison, statistical analysis was performed using Welch’s t-test.

## Acknowledgements

Schematics were created in https://BioRender.com. The research reported in this publication was supported by the University of Michigan Advanced Genomics Core, the UM Single Cell Spatial Analysis Program and the National Cancer Institutes of Health under Award Number P30CA046592 using the following Cancer Center Shared Resource: Single Cell and Spatial Analysis Shared Resource. We would also like to thank the University of Michigan Flow Cytometry Core, especially Dr. Sylvaine Lambert and Dr. Andrea Hodgins-Davis, for technical assistance and troubleshooting of the sorting and flow analysis experiments. We thank Dr.

Aaron Morris for help with the data curation for the EAE model. We thank Sargis Shameon from the Lyssiotis Lab for help with peak annotation for the LC-MS/MS data. We would also like to thank other members of the Shea Lab, especially Dr. Russell Urie and Dr. Sean Carey for assisting with various experimental and analysis troubleshooting.

## Supplementary information

The supplementary files related to blood glucose, gene expression analysis, metabolomics analysis are available in Supplementary Tables 1-10 and Supplementary Figures 1-8.

## Data availability

The IN bulk RNA sequencing from NOD and pancreas single cell RNA sequencing data will be available in GEO upon acceptance. A part of the NOD IN bulk RNA sequencing data was taken from GEO: GSE299709. The 4T1 breast cancer data IN bulk RNA sequencing data was obtained from GEO: GSE233001. The adoptive transfer T1D transcriptomic data was obtained from King *et al*.(20).

HPAP scMultiome data from spleen and pancreatic lymph node (PLN) were downloaded from PanCDB under the HPAP user agreement. Access to these datasets is available through the HPAP portal (https://hpap.pmacs.upenn.edu/) following HPAP data access procedures. This manuscript used data acquired from the database (https://hpap.pmacs.upenn.edu/) of the Human Pancreas Analysis Program (HPAP; RRID:SCR_016202; PMID: 31127054; PMID: 36206763). HPAP is part of a Human Islet Research Network (RRID:SCR_014393) consortium (UC4-DK112217, U01-DK123594, UC4-DK112232, and U01-DK123716). PanKbase annotated

V3.3 HPAP single cell expression map of pancreatic islets was downloaded from PanKbase under the PanKbase user agreement. Access to these datasets is available through the PanKbase portal (https://pankbase.org/). PanKbase (https://www.pankbase.org/), supported by NIDDK U24DK138515, U24DK138512, and supplemental funds from the NIH Office of Data Science Strategies.

All the flow data will be available in the Deep Blue Repositories University of Michigan Library after acceptance.

## Author contributions

Conceptualization: J.R. and L.D.S. Methodology: J.R., Y.J., R.M., J.L.K, A.T., L.H., H.Z., X.L., P.S., B.H., K.L, J.L, C.A.L., and L.D.S. Software: J.R., Y.J., R.M., A.T., H.Z., X.L and K.L. Investigation: J.R., Y.J., R.M. and L.D.S. Formal analysis: J.R., Y.J., R.M., J.L.K, A.T., and P.S. Data curation: J.R., Y.J., R.M, J.L.K., P.S., B.H., E.J.B., L.M.R., S.S., A.H., B.W, K.K., D.A., and A.H.M. Writing—original draft: J.R. and L.D.S. Writing—review and editing: J.R., Y.J., R.M., P.S., E.J.B., D.A., and L.D.S. Visualization: J.R., Y.J. and L.D.S. Supervision: L.D.S., J.L., and C.A.L. Project administration: J.R. and L.D.S. Funding acquisition: J.R. and L.D.S.

## Code availability

Codes used in this study is publicly available in Github: https://github.com/shea-lab/IN_T1D_Immune_TNF_Paper

