## Supplementary Figures for "Synthetic Immunological Niche Reveals Early Immune Dysregulation and Stratifies Therapeutic Response in Type 1 Diabetes"

### a Myeloid Gating

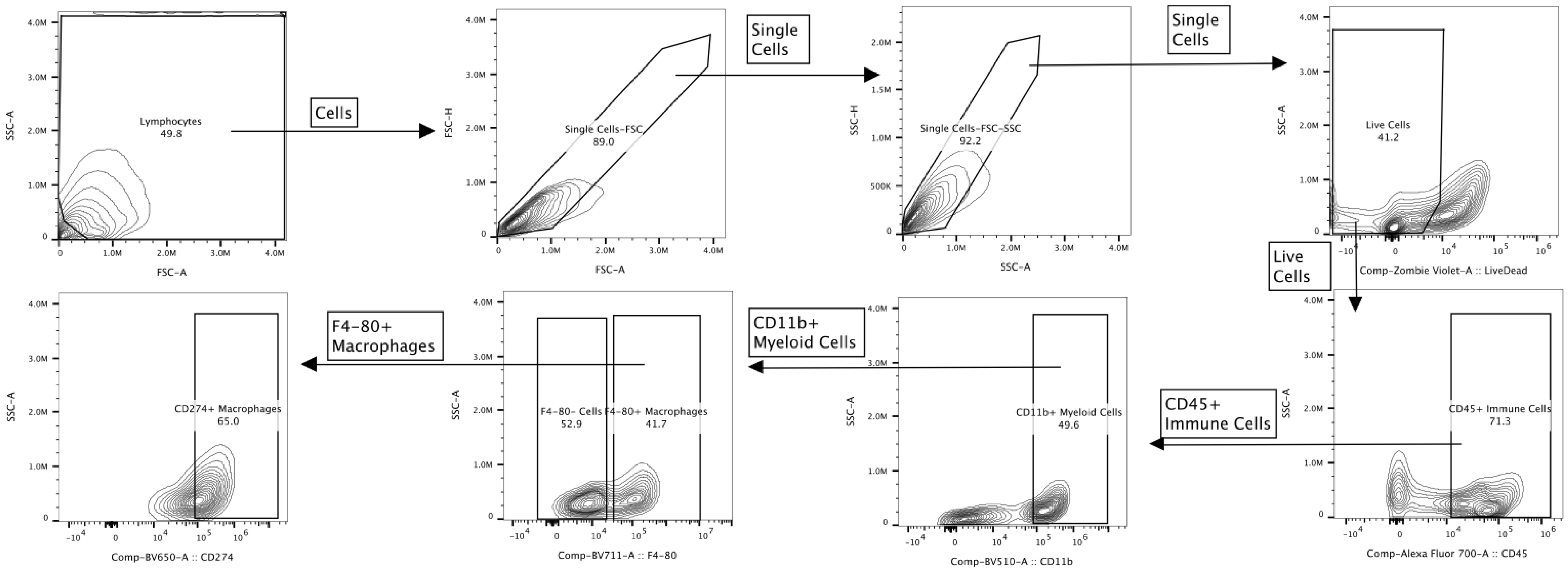


### b Lymphoid Gating

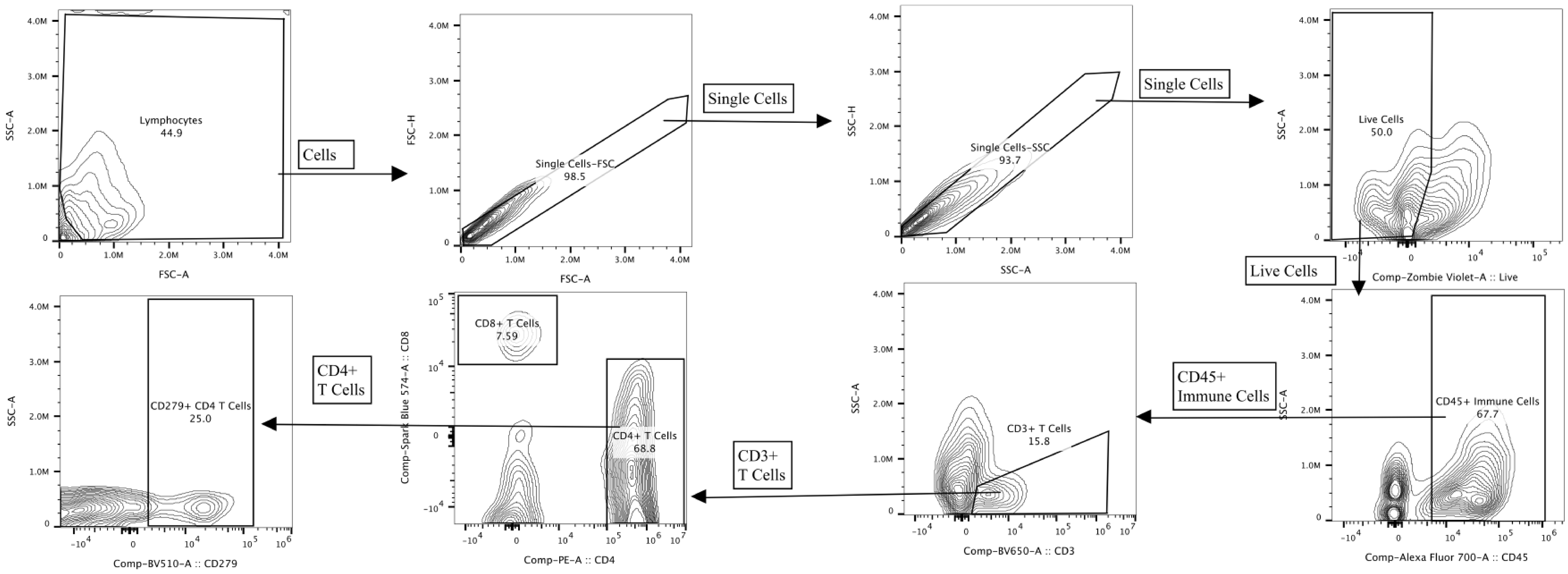


**Figure S1**: **Flow Cytometry Gating Strategy-** Flow cytometry gating strategy for **a)** Myeloid and **b)** Lymphoid panel

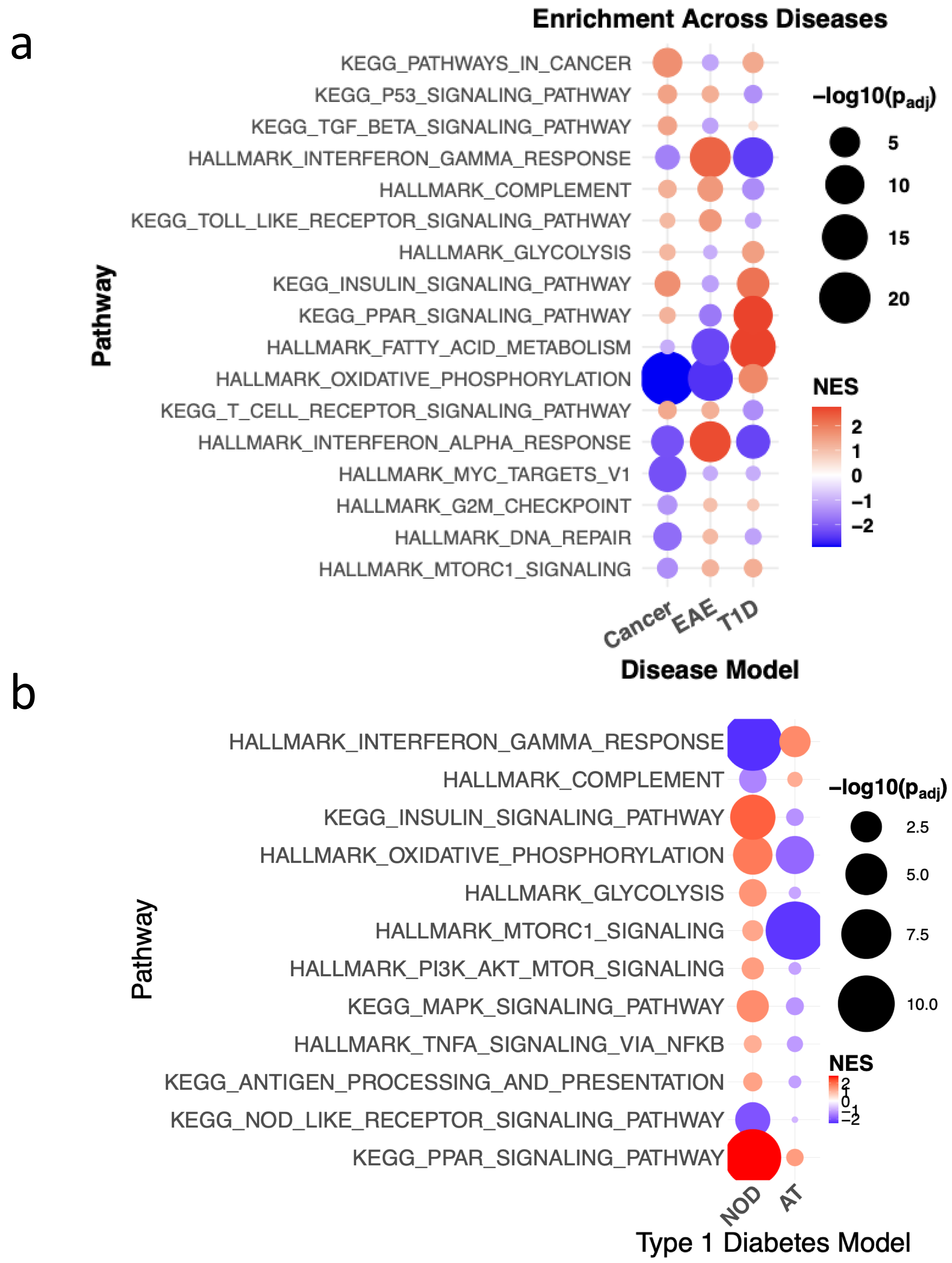


**Figure S2**: **Geneset Enrichment Analysis of IN transcriptomics-** Dotplot showing Normalized Enrichment Score and significance for selected pathways for- **a)** Early Stage NOD T1D vs 4T1 Breast Cancer and Experimental Autoimmune Encephalomyelitis. inflammation disease comparison **b)** Early Stage NOD vs Adoptive Transfer (AT) T1D models

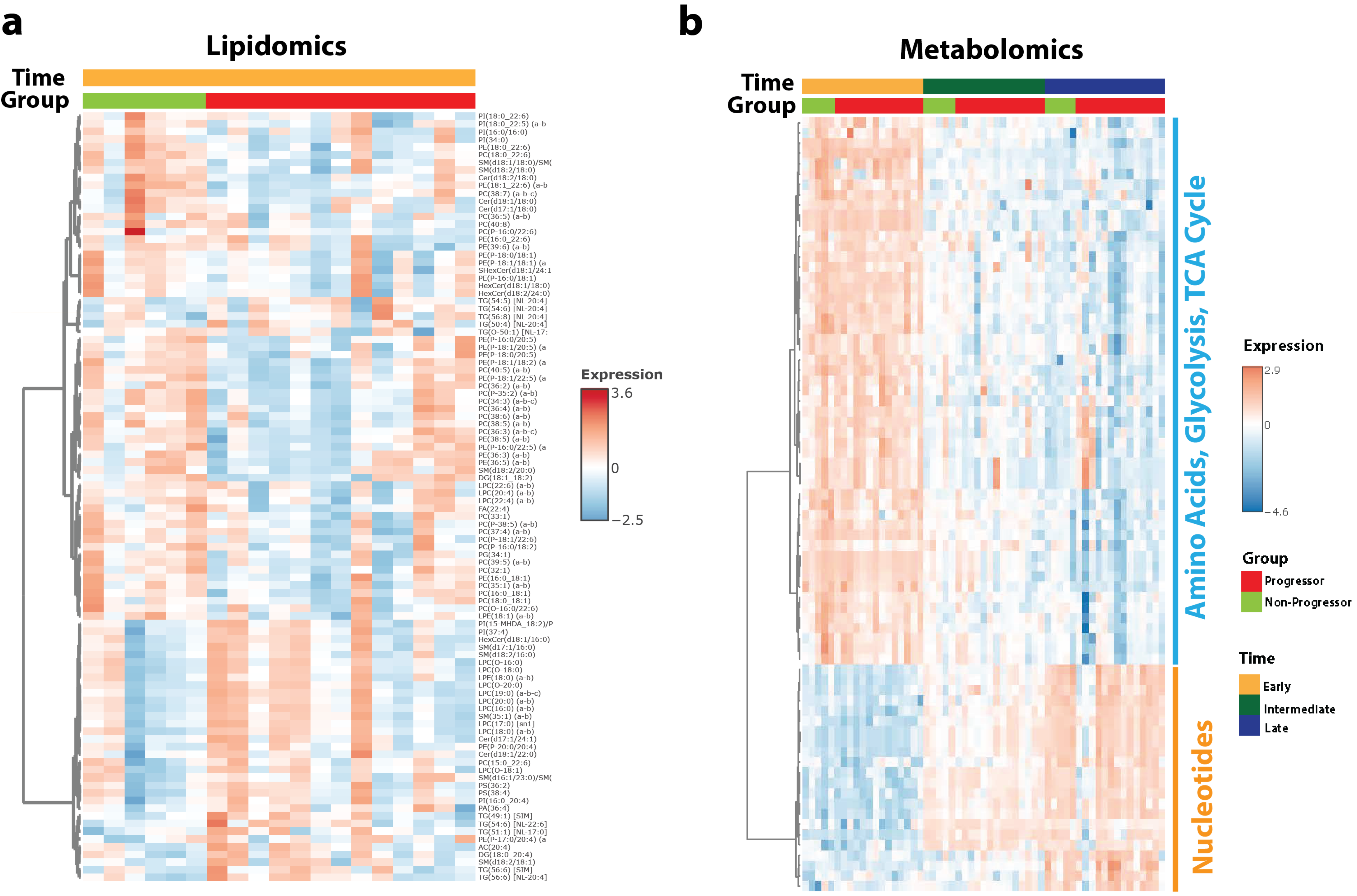


**Figure S3**: **Lipidomic and Metabolomic profiling of the Immunological Niche- a)** Lipid profiles at the early stage between progressor(n=13) vs non-progressors (n=6) **b)** Polar metabolite profiles at the different T1D stages between progressor(n=14) vs non-progressors (n=4)

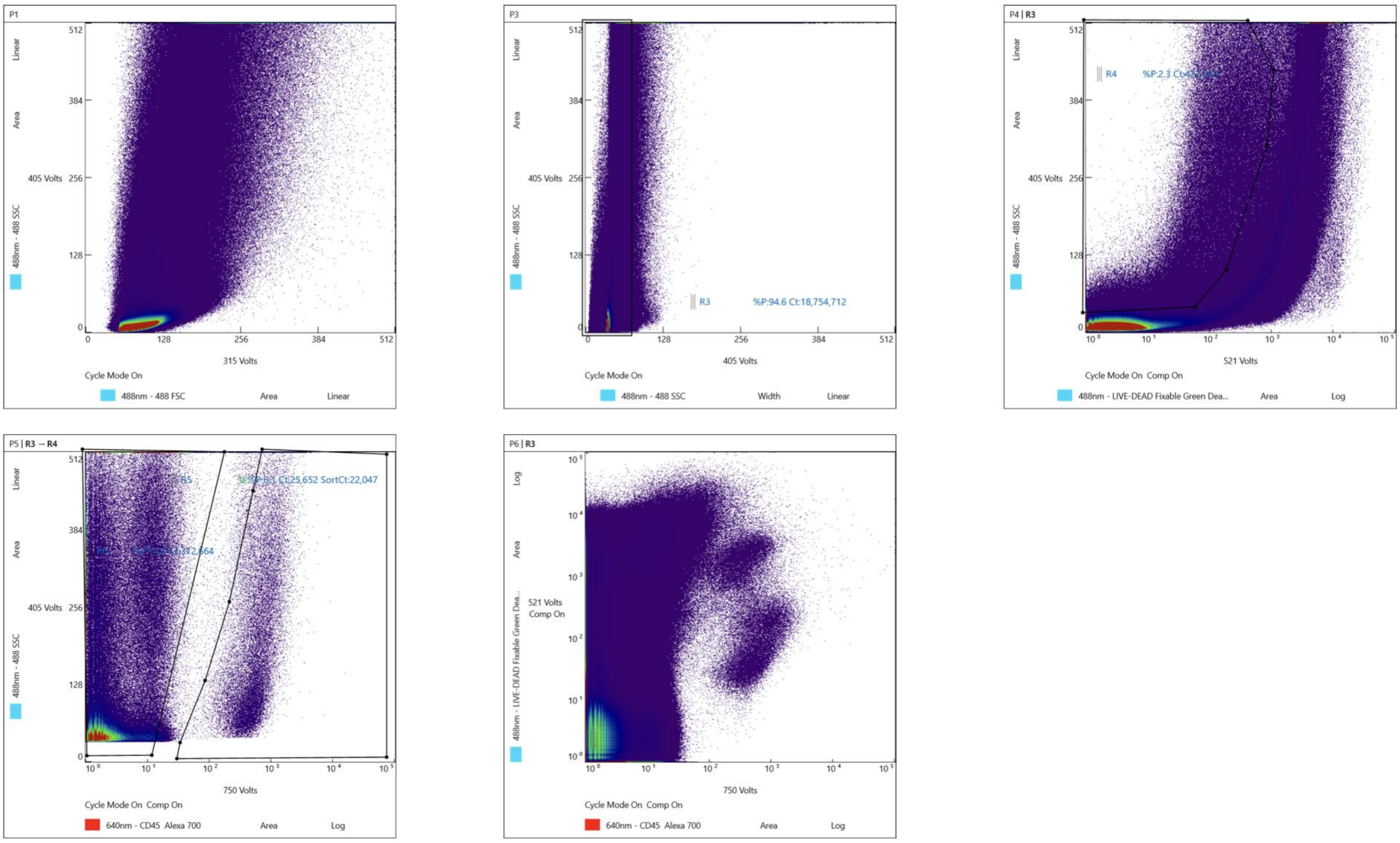


**Figure S4**: **Gating Strategy For NOD Pancreas Immune Cell Sorting using FACS-** Immune cells were sorted based of Live CD45+ gating

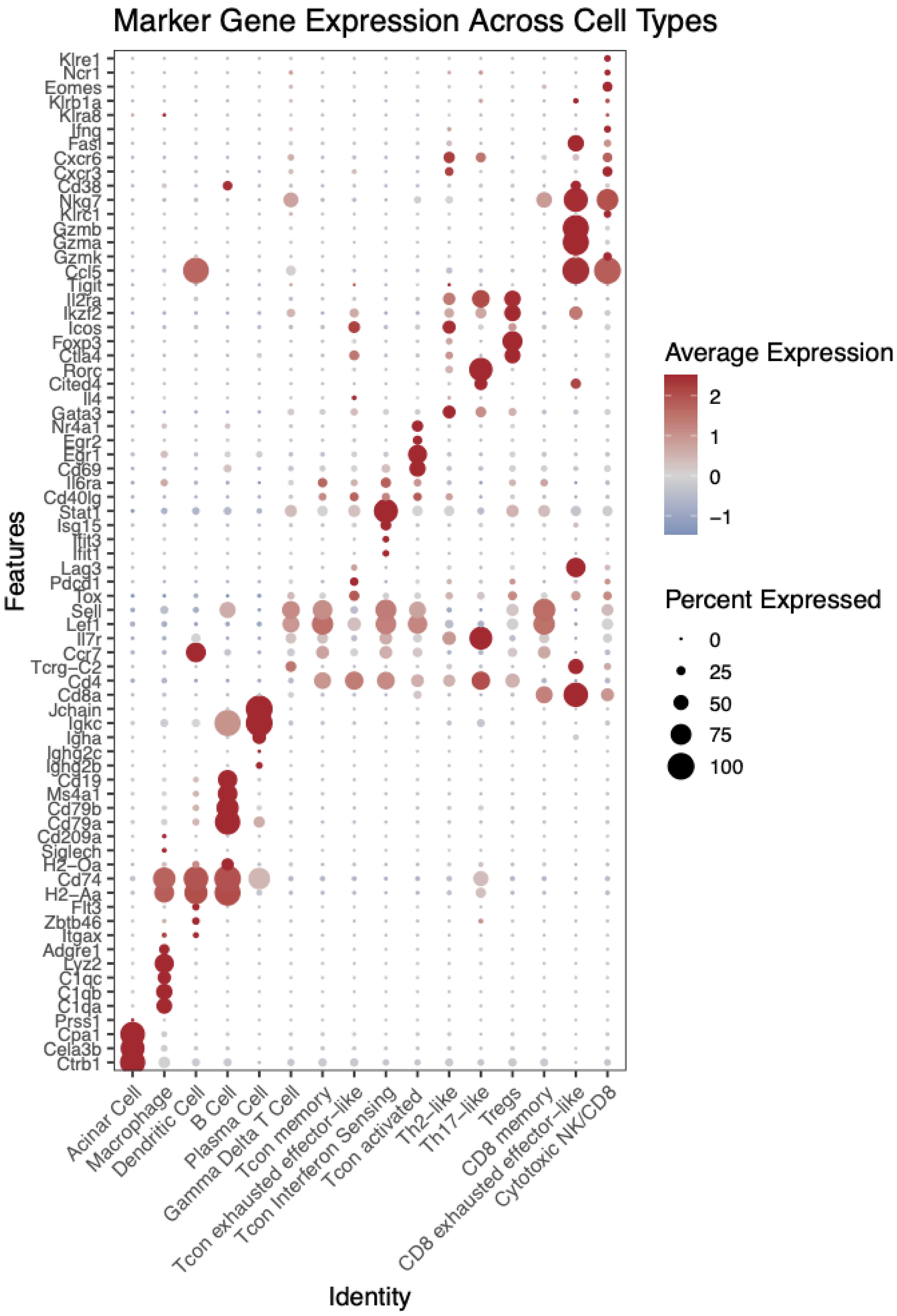


**Fig. S5**. **Dot plot of canonical marker genes used for annotation of pancreas cell types-** Scaled expression levels and frequency of canonical genes are shown for each identified NOD pancreatic cells

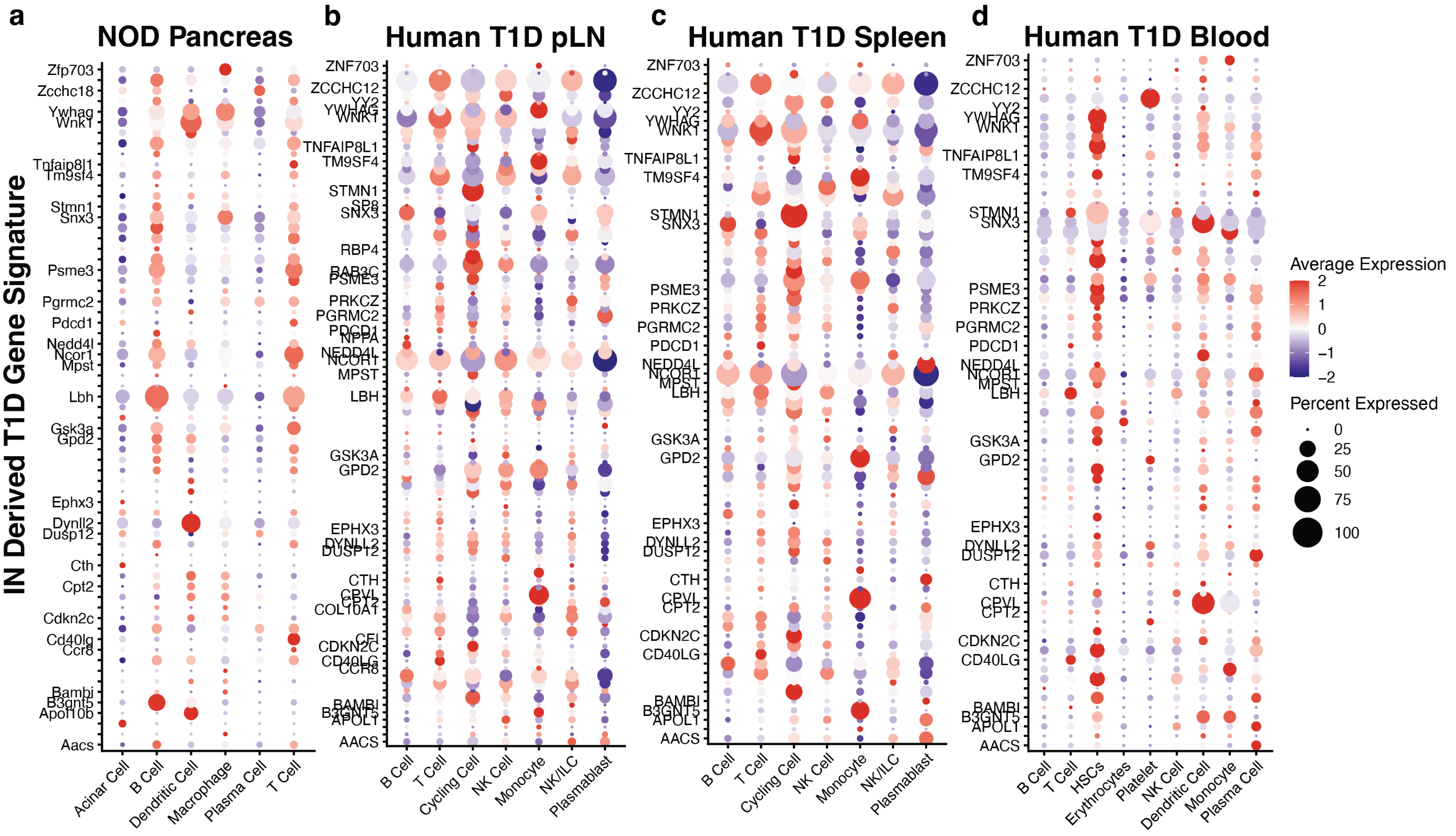


**Fig. S6**. **Mapping of IN derived T1D gene signature.** Scaled expression levels and frequency of T1D gene signature (or their human orthologs) for **a)** NOD Pancreas, and **b)** Human pLN, **c)** spleen, **d)** PBMCs from T1D and non-diabetic individuals

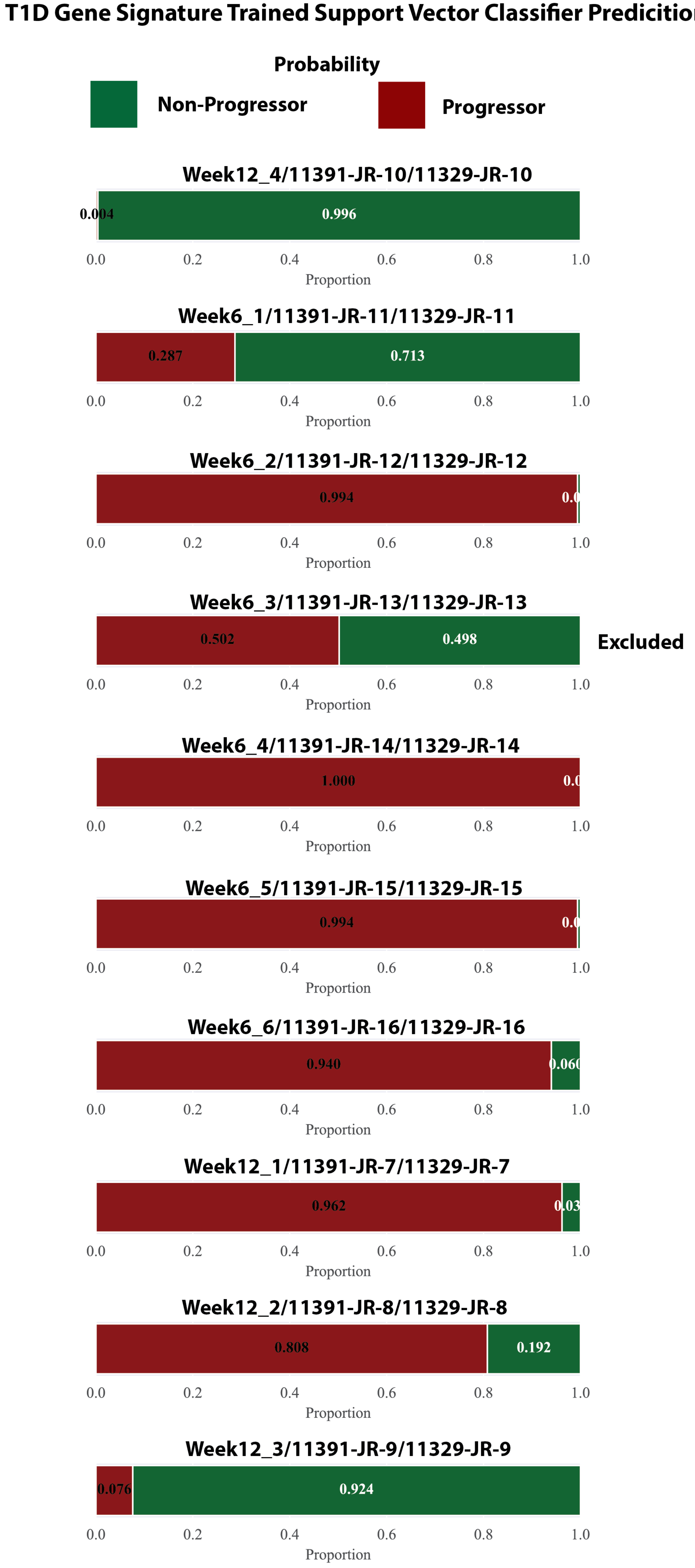


WeekA_B- Mouse Number For ScRNA Sequencing (A-Time, B-Mouse)
11391-JR-X- Sample ID For IN Bulk RNA Sequencing
11329-JR-X- Sample ID For Pancreas ScRNA Sequencing

**Fig. S7. Prediction of NOD as Progressor vs Non-Progressor Using IN Derived T1D 100 Gene Signature.** Probability of being Progressor as predicted by the Support Vector Classifier trained on IN Derived T1D 100 Gene Signature. Samples classified with >60% probability were taken for downstream analysis of the pancreas single cell RNA sequencing data
