## Supplementary Methods for "Synthetic Immunological Niche Reveals Early Immune Dysregulation and Stratifies Therapeutic Response in Type 1 Diabetes"

**Methods Supplementary File**

*Human Tissue Single Cell Pre-Processing*

We analyzed HPAP spleen and pancreatic lymph node (PLN) snMultiome (RNA+ATAC) datasets (1); no standalone scRNA-seq datasets were available for these tissues. Downstream analyses in this study (dimensional reduction, clustering, cell-type annotation, and differential expression) were performed using the RNA modality only. However, because the raw data were generated with the snMultiome protocol, nucleus identification and quality control were performed using joint RNA and ATAC metrics to mitigate modality-specific artifacts and ensure consistent filtering across heterogeneous donors and processing centers. Accordingly, we describe ATAC preprocessing and ATAC-derived QC metrics only to the extent that they informed nucleus-level QC and filtering.

RNA reads were aligned and quantified at the donor level, followed by barcode calling and decontamination. Putative cell-containing barcodes were identified using emptyDrops from DropletUtils (2). To mitigate ambient RNA contamination, CellBender “remove-background” outputs were loaded from the CellBender H5, retaining inferred latent variables. In the joint QC stage, we used CellBender-derived quantities such as cell probability and per-barcode post-decontamination UMI counts to compute the fraction of ambient signal removed and to support RNA-side filtering (3).

ATAC reads were adapter-trimmed (CTA), assessed with FastQC/MultiQC, and aligned to the reference genome using BWA-MEM with coordinate sorting by SAMtools (4–6). Read groups belonging to the same library were merged, duplicates were marked with Picard, and a high-confidence ATAC alignment set was obtained by retaining properly paired autosomal reads with MAPQ ≥ 30 while removing unmapped, secondary, duplicate, and supplementary alignments.

Broad ATAC peaks were called with MACS2, and these peaks together with the processed BAMs were used as inputs to ataqv to compute ATAC QC summaries, including TSS enrichment, mitochondrial read fraction, and high-quality autosomal accessibility (HQAA) as well as the data-driven suggested HQAA cutoff reported by ataqv, enabling sample-adaptive ATAC quality filtering across heterogeneous donors/centers (7, 8).

RNA and ATAC metrics were then integrated per nucleus, producing a single metrics table per donor that retained both modalities (RNA UMI depth, RNA %mitochondrial, CellBender-derived metrics; ATAC HQAA, ATAC TSS enrichment, ATAC %mitochondrial). Because HPAP snMultiome data are not uniformly generated across donors and centers, we avoided fixed global cutoffs and instead estimated several key thresholds per donor using data-driven procedures. In particular, we computed modality-aware mitochondrial thresholds by first diagnosing whether the underlying distribution was unimodal or multimodal via KDE peak counting, then applying either 2D histogram smoothing and Multi-Otsu segmentation to separate foreground high-quality nuclei from background in the joint space of accessibility/UMIs vs. %mitochondrial, or fallback 1D Multi-Otsu thresholding when the 2D segmentation was unstable. This same philosophy was used to set additional RNA-side cutoffs using Multi-Otsu on the observed distributions. We also performed a refined knee-plot analysis of barcode-rank vs. RNA UMI curves, including interpolation on a log scale and Savitzky–Golay smoothing to stabilize slope-based cliff detection in noisy donors.

After applying the RNA- and ATAC-side preprocessing and the joint, per-donor QC filtering, doublets were annotated on the QC-passing snMultiome nuclei using two complementary strategies to reduce modality-specific artifacts: AMULET and DoubletFinder (9). Concordance between the two calls was used to summarize doublet burden and to support downstream cluster-level artifact screening. The filtered nuclei were then processed in Seurat using normalization, variable feature selection, PCA with PC selection guided by elbow plots, and initial UMAP clustering to establish baseline structure prior to correction. To mitigate non-biological structure driven by donor and processing covariates, we applied Harmony on the PCA space using covariates including HPAP donor ID, sex, BMI, age, ethnicity, and relocation center, producing corrected embeddings. Integration performance was assessed by comparing pre- and post-Harmony diagnostics, with a primary criterion that no cluster was dominated by a single metadata category, while preserving expected biological marker patterns (10). Finally, low-quality or artifactual clusters were flagged for removal based on high doublet proportion together with incoherent marker expression, and by abnormal cluster-level QC distributions defined as follows: for %chrMT, clusters were flagged when the fold change < 1/2 and the associated adjusted p-value < 0.05; for nUMI and nFeature, clusters were flagged when the fold change > 2 with an adjusted p-value < 0.05. For cell annotation, we leveraged an externally curated HPAP CITE-seq reference dataset with manual cell-type annotations as the reference panel. Cluster-level reference mapping was performed using SingleR to get the final annotation (11). Cells were called restricted to donors annotated as non-diabetic (ND) or stage 3 type 1 diabetes (T1D) based on annotation from PanKbase (12). Harmony-derived clusters were mapped to major immune cell types.

*Metabolomics and Lipidomics Analysis*

For targeted metabolomics, the immunological niche was bead-homogenized, and metabolites were extracted using 80% methanol (v/v). Supernatants from each sample were collected based on weight normalization. The collected supernatants were then dried using a SpeedVac Vacuum Concentrator, reconstituted in 50% methanol (v/v) in water, Targeted metabolomics was performed on an Agilent 1290 Infinity II Binary Bio LC coupled with an Agilent 6495d QqQ mass spectrometer. The column used was an Agilent InfinityLab Poroshell 120 HILIC-Z, 2.1 x 150 mm, 2.7uM (p/n 683775-924). Method parameters are as follows: Solvent A is water +

20mM ammonium acetate (pH 9.3) + 5uM medronic acid. Solvent B is acetonitrile. Wash solvent is 1:1:1 water, acetonitrile, and methanol. The solvent gradient is 10% A – 90% B until 1 minute, 22% A – 78% B until 8 minutes, 40% A – 60% B until 12 minutes, 90% A – 10% B until 15 minutes, hold until 18 minutes, 10% A – 90% B until 19 minutes, hold until 23 minutes. Flow rate is 0.4mL/minutes. Column temperature is set to 15 C. Source parameters are optimized for the Agilent dMRM (dynamic multiple reaction monitoring) library. The dMRM library was acquired from Agilent for their HILIC-Z platform and further optimized to screen for 435 targets in both positive and negative ion modes.

Lipids were extracted from the immunological niche using a chilled biphasic solvent system (MTBE:methanol, 10:3, v/v). Samples were homogenized, phase-separated with water, and the upper organic phase was collected. Targeted lipidomics was performed on an Agilent 1290 Infinity II Binary Bio LC coupled with an Agilent 6495d QqQ mass spectrometer. The column used was an Agilent ZORBAX Eclipse Plus C18, 100 × 2.1 mm, 1.8 µm (p/n 959758-902). Method parameters are as follows: Solvent A consists of 10 mM ammonium formate and 5 µM Agilent deactivator additive (p/n 5191-3940) in 5:3:2 water: acetonitrile:2-propanol. Solvent B consists of 10 mM ammonium formate in 1:9:90 water: acetonitrile:2-propanol. Wash solvent is a 1:1 mixture of solvents A and B. The solvent gradient is 85% A – 15% B until 2.5 minutes, 50% A – 50% B until 2.6 minutes, 43% A – 57% B until 9 minutes, 30% A – 70% B until 9.1 minutes, 7% A – 93% B until 11 minutes, 4% A – 96% B until 11.1 minutes 0% A – 100% B until 12.2, 85% A –15% B until 16 minutes. Flow rate is 0.4mL/minutes. Column temperature is set to 45 C. Samples are kept at 20 C. The dMRM library was acquired from Agilent for targeted lipidomics platform and further optimized to screen for 665 targets in both positive and negative ion modes.

Data from both methods was preprocessed using Agilent MassHunter Workstation Quantitative Analysis for QQQ Version 12.1. 12.1. Raw data was normalized by median-centering and pareto normalization then further fold-change and Student's T- Tests analysis was performed relative to control groups to achieve a relative abundance in Metaboanalyst (v. 5.0).
