## Supplementary Tables 1-10 for "Synthetic Immunological Niche Reveals Early Immune Dysregulation and Stratifies Therapeutic Response in Type 1 Diabetes": SupplementaryTableLegends.docx

**Supplementary Tables Description**

**Supplementary Table 1 (ST1). Cohort Characterization and Metabolic Phenotyping**
ST1 contains longitudinal blood glucose measurements for all experimental cohorts, including weekly glucose monitoring data. It also includes intraperitoneal glucose tolerance test (IPGTT) results.

**Supplementary Table 2 (ST2). Flow Cytometry Immune Profiling**
ST2 provides flow cytometry—based immune cell proportions quantified from the IN for the myeloid and lymphoid compartments. Data is reported as frequencies of total CD45⁺ cells for each time point and mouse.

**Supplementary Table 3 (ST3). NOD Progressor vs Non-Progressor IN Transcriptomic Analysis**
ST3 contains differential gene expression (DEG) results comparing IN in T1D progressors and non-progressors at each stage- early, intermediate and late. The table also includes stage-wise Gene Set Enrichment Analysis (GSEA) results.

**Supplementary Table 4 (ST4). IN T1D Specificity Analysis**
ST4 presents differential expression and pathway analyses used to assess T1D specificity of the IN-derived early stage DEGs. This includes DEG lists from 4T1 Breast cancer, Experimental Autoimmune Encephalomyelitis (EAE), and adoptive transfer (AT) T1D models, along with corresponding GSEA-enriched pathways for each model. Diseased vs healthy was compared in each model

**Supplementary Table 5 (ST5). Metabolomics and Lipidomics Profiling**
ST5 includes metabolomic and lipidomic profiling of the IN microenvironment for progressors vs non-progressors. The lipidomics data is for early timepoint and the metabolomics data is across all time points.

**Supplementary Table 6 (ST6). PLS-DA Modeling Results**
ST6 has Partial Least Squares Discriminant Analysis (PLS-DA) results derived from IN transcriptomic data at early stages. It includes the top 100 genes ranked by Variable Importance in Projection (VIP) scores, as well as scaled expression values used for heatmap visualization.

**Supplementary Table 7 (ST7). Macrophage-Specific Progressor vs Non-Progressor Analysis**
ST7 provides differential gene expression results from pancreas macrophage populations, comparing T1D progressors and non-progressor NODs.

**Supplementary Table 8 (ST8). IN-Based Incidence and Anti-TNF Cohort Data**
ST8 contains longitudinal blood glucose measurements for anti-TNF-treated cohorts and untreated controls.

**Supplementary Table 9 (ST9). Anti-TNF Sensitive vs Resistant IN Transcriptomic Analysis**
ST9 provides differential gene expression of IN comparing anti-TNF sensitive and resistant groups. It includes pre-treatment and post-treatment DEG analyses, as well as GSEA-enriched pathways for each condition.

**Supplementary Table 10 (ST10). IN-Derived Anti-TNF Response Score (ATRS)**
ST10 contains input gene sets related to TNF for elastic net, obtained parameters used for calculating the ATRS for selected gene sets, as well as individual ATRS scores for each sample.
